# Mucosal B cell memory selection integrates tissue-specific cues via the IgA BCR

**DOI:** 10.1101/2025.04.30.651421

**Authors:** Maria Pia Holgado, Samuel Origlio, Lucas Bertoia, Myriam Moussa, Apurwa Trivedi, Frederic Fiore, Ana Zarubica, Emma Slack, Michelle A. Linterman, Michel Cogné, Claude Gregoire, Mauro Gaya

## Abstract

B cells engender plasma cells (PCs) and memory B cells (MBCs) to combat recurrent pathogens. While B cell receptor (BCR) affinity dictates fate decisions in clonally-restricted set-ups, the driving factors in more complex systems remain unknown. Here, we show that memory selection during mucosal infection is imprinted by inherent cues from barrier tissues: lungs skew selection towards MBCs while the gut favors PC entry, even in response to the same pathogen. This divergence is linked to differential BCR isotype usage across barrier tissues rather than to affinity. In the gut, the commensal-induced TGF-β-rich milieu promotes B cell class-switching to IgA. Consequently, the IgA BCR bias selection towards PCs, a process that is counteracted by its cytosolic tail domain. Overall, mucosal B cell selection integrates tissue-specific cues by strongly relying on BCR isotype usage rather than affinity, with implications for nasal and oral vaccine development.

**One sentence summary:** Divergent B cell memory strategies evolved in lung and gut mucosa to counteract recurrent pathogens.

## Introduction

One of the main facets of vertebrates’ immune system is the ability to remember previously encountered pathogens through a process known as immunological memory. This strategy involves the generation of long-lasting memory cell populations derived from B and T lymphocytes. In particular, B cells can give rise to two different memory compartments: plasma cells (PCs), which constitutively secrete antibodies throughout life, and memory B cells (MBCs), which remain in quiescent state but can rapidly differentiate into antibody-secreting cells upon antigen re-exposure, complementing existing humoral immunity (*1*). MBCs and PCs are the basis of vaccination, therefore, deciphering the cellular and molecular mechanisms that drive B cell selection into the PC and MBC programs is of utmost interest.

Previous work using immunization with model antigens or clonally restricted B cell systems, showed that BCR affinity plays a key role in determining memory cell fate: high affinity B cells preferentially differentiate into PCs while the MBC compartment is less stringent, allowing for the entry of low-affinity cells and supporting memory diversity (*2–4*). Recent work revealed that immunization with complex antigens supports the output of low- and high affinity PCs, suggesting that affinity is not the only factor driving the PC program when multiple B cell clones with distinct BCR specificities coexist (*5*, *6*).

Unlike systemic immunization with model antigens, which predominantly induces IgM- and IgG-expressing B cells with limited IgA responses, mucosal infection gives rise to B cells with diverse specificities and BCR isotypes (IgM, IgG, and IgA) that take up residence not only in lymphoid organs but also in barrier tissues such as the lungs (*7–11*) and gut (*12–16*). This raises the question of the mechanisms that dictate B cell memory selection in such scenarios. Here, we used respiratory and gastrointestinal infection models to investigate the mechanisms instructing B cell memory in the mucosas. We found that inherent features from barrier tissues alternately skews B cell selection towards MBCs in the lungs and PCs in the gut. Limiting affinity maturation by means of loss-of-function mutation in Aid did not impact memory cell fate. Instead, differential BCR isotype expression across barrier tissues accounts for the tissue-specific biased selection. We found that preferential usage of IgA BCRs in the gut, fostered by a commensal-induced TGF-β-rich milieu, fine-tunes downstream signaling upon antigen encounter and favours B cell entry into the PC program. Then, we propose a model where mucosal B cell memory selection integrates tissue-specific cues by strongly relying on BCR isotype usage rather than affinity.

## Results

### Divergent B cell memory strategies evolved in lungs and gut

To study B cell memory responses in the mucosa, we used the *Aicda-*Cre^ERT2^ *Rosa26*-EYFP (Aid-EYFP) mouse model, which allows permanent labeling of PCs and MBCs based on their precursors’ expression of *Aicda*, the enzyme that initiates class-switch recombination and somatic hypermutation (*17*). We initially analyzed the establishment of B cell memory in the lungs and gut, as these organs are key barrier tissues exposed to the environment and play a central role in orchestrating homeostatic and pathogenic immune responses.

To analyze lung B cell memory, Aid-EYFP mice were intranasally exposed to PBS or influenza A virus PR8 strain, followed by tamoxifen treatment beginning on day 6 (Figure 1A). After 5 weeks, mice were intravenously injected with labeled anti-CD45 antibody to exclude cells from the blood circulation from our analysis. Lungs and mediastinal lymph nodes (med LNs) were then harvested. We found that influenza infection triggered the accumulation of YFP^+^ cells in both organs, whereas YFP⁺ cells remained negligible in PBS-treated mice (Figures 1B and S1A). Resident YFP^+^ cells were then segregated into memory and germinal center (GC) B cells based on GL7 expression by flow cytometry (Figures 1C and S1B). GC B cells were detected in both the lungs and med LNs. Within the memory compartment, PCs and MBCs were present at an approximate 25:75 ratio, resulting in a PC/MBC ratio significantly lower than 1 and indicating that the lung B cell memory compartment is enriched in MBCs (Figures 1C-1D and S1C). A similar trend was observed in the med LNs (Figure S1B). Both PCs and MBCs recognized influenza hemagglutinin (HA) and nucleoprotein (NP) antigens but localized to distinct lung regions: PCs formed clusters at the periphery of bronchus-associated lymphoid tissue, whereas MBCs accumulated in the center of these tertiary lymphoid structures (Figures 1E-1G). To assess the stability of this MBC-biased composition, mice were treated as in Figure 1A, tamoxifen administration was stopped at week 5, and animals were sacrificed 365 days after infection. MBCs largely outnumbered PCs in the lung even one year post-infection, demonstrating that this MBC-biased organization is maintained over time once established (Figures S1D-S1E). In addition, this experiment confirms that lung MBCs and PCs derive from cells engaged early during the response, as they were labeled by tamoxifen during the first month of infection and were still detected in the lungs 11 months later.

**Figure 1.**
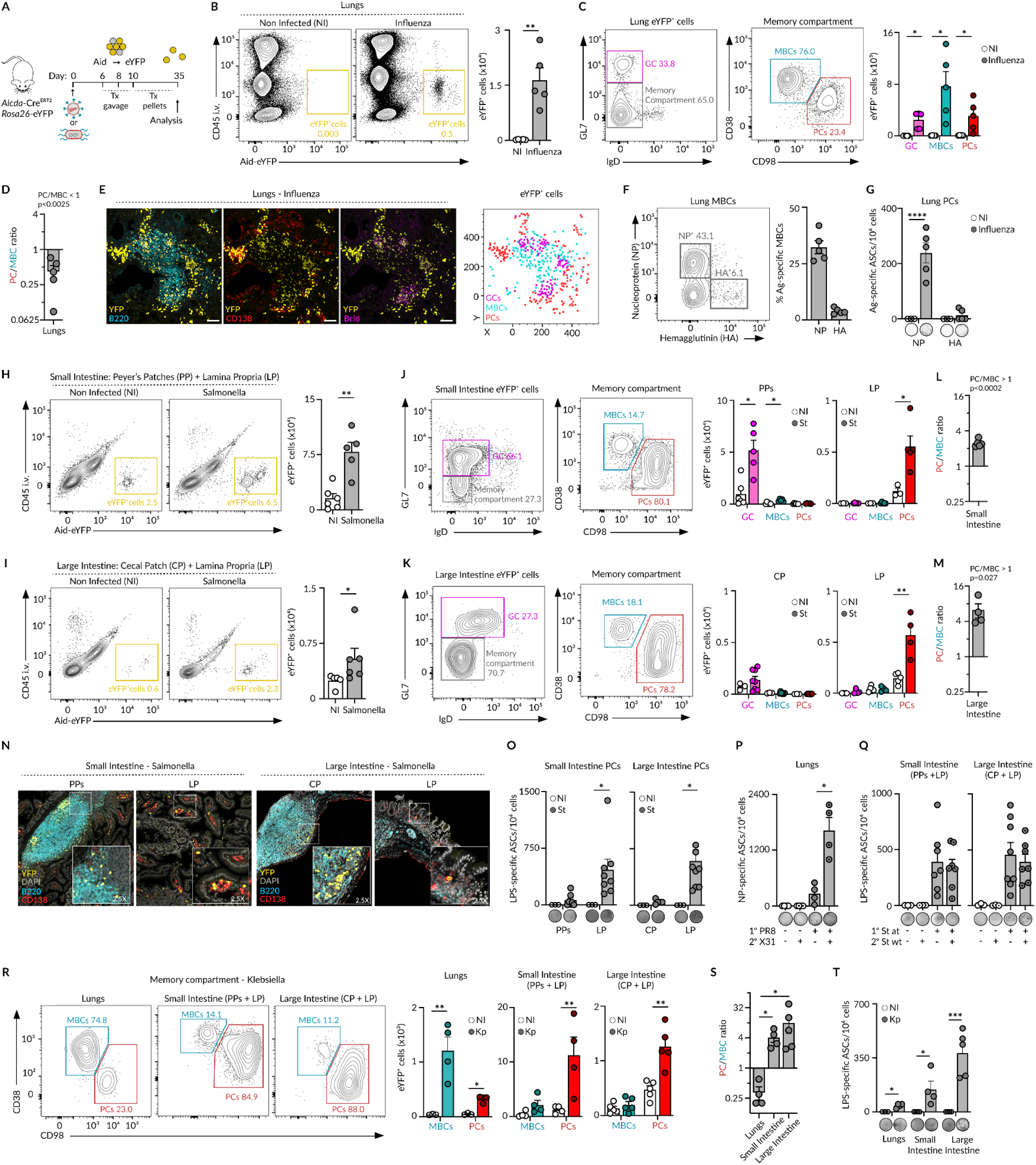
Mucosal infection elicits divergent B cell memory strategies in lungs and gut. (**A**) Overview of experimental approach. (**B**) Flow cytometry plots of total lung cells in non-infected (NI) or influenza-infected AID-EYFP mice treated as in (A). (**C**) Flow cytometry plots displaying the gating strategy of lung YFP^+^ GC B cells (GL7^+^), memory cells (GL7^-^), MBCs (CD38^+^CD98^-^), and PCs (CD38^low^CD98^+^) in Aid-EYFP mice infected with influenza virus and treated as in (A). Quantifications indicate absolute numbers of total eYFP+ cells (B) or GC B cells, MBCs and PCs (C). (D) Bar chart showing the PC:MBC ratio in lungs of mice infected with influenza virus. (**E**) Confocal images of lung sections stained for YFP (yellow), B220 (cyan), CD138 (red), and Bcl6 (magenta). Scale bars, 70 μm. X and Y positions of MBCs (B220^+^Bcl6^-^, cyan), GCs (B220^+^Bcl6^+^, magenta) and PCs (B220^low^CD138^+^, red). (**F**) Contour plot and quantification of lung YFP^+^ MBCs binding to NP and HA antigens. (**G**) Enumeration of lung NP- and HA-specific antibody-secreting cells by ELISpot. (**H** and **I**) Flow cytometry plots of total gut YFP^+^ B cells in (H) small (Peyer’s patches and lamina propria) and (K) large (cecal patch and lamina propria) intestine in non-infected or *Salmonella typhimurium -* infected mice treated as in (A). (**J** and **K**) Flow cytometry plots displaying the gating strategy of (J) small and (K) large intestine YFP^+^ GC B cells (GL7^+^), memory cells (GL7^-^), MBCs (CD38^+^CD98^-^), and PCs (CD38^low^CD98^+^) in Aid-EYFP mice infected with *Salmonella typhimurium* and treated as in (A). Quantifications indicate absolute numbers of total eYFP+ cells (H and I) or GC B cells, MBCs and PCs (J and K). (**L** and **M**) Bar charts showing the PC:MBC ratio in (L) small or (M) large intestine of mice infected with *Salmonella typhimurium*. (**N**) Confocal microscopy images of small (Peyer’s patches and lamina propria) and large (cecal patch and lamina propria) intestine sections stained for YFP (yellow), B220 (cyan), CD138 (red) and DAPI (grey). (**O**) Enumeration of intestinal Salmonella LPS-specific antibody-secreting cells by ELISpot. (**P** and **Q**) Enumeration of (P) NP-specific or (Q) LPS-specific antibody-secreting cells in lungs or intestine after re-challenge with influenza X-31 or *Salmonella* wild type, respectively. For (Q), pooled data from two experiments is shown. (**R**) Flow cytometry analysis of MBCs and PCs in lungs and intestine of Aid-EYFP mice following *Klebsiella pneumoniae* infection. Quantifications indicate absolute numbers of eYFP+ MBCs and PCs (**S**) Bar chart showing the PC:MBC ratio in lungs and intestine of mice infected with *Klebsiella pneumoniae* by intranasal or oral infection, respectively. (**T**) Enumeration of Klebsiella LPS-specific antibody-secreting cells by ELISpot. In all panels, unless indicated, quantifications display one representative experiment out of three, mean ± s.e.m. Each dot represents one mouse. For statistical analysis, we used t-test (B, H, I), one-way Anova (S) or two-way Anova (C, G, J, K, O, P, Q, R, T). For panels (D, L and M) we perform a t-test to calculate whether the PC/MBC ratio was < or > 1. *p<0.05, **p<0.01, ***p<0.001 and ****p<0.0001.

To determine the composition of the B cell memory compartment in the gut mucosa, Aid-EYFP mice were orally administered PBS or *Salmonella typhimurium* strain SL3261, followed by tamoxifen treatment beginning on day 6 (Figure 1A). After 5 weeks, the small intestine (Peyer’s patches and lamina propria), large intestine (cecal patch and lamina propria), and mesenteric (mes) LNs were harvested. As expected, we detected baseline levels of YFP⁺ cells in all organs of PBS-treated mice, reflecting ongoing immune responses to the gut microbiota (Figures 1H-1I and S1F). Following Salmonella infection, we observed a robust and significant increase in YFP⁺ cells in Peyer’s patches, lamina propria, and mes LNs (Figures 1H-1K and S1F-S1G). In contrast, YFP⁺ cell numbers in the cecal patch did not increase above baseline levels, consistent with the preferential targeting of Peyer’s patches during Salmonella infection (Figure 1K). Together, these data indicate that although the YFP⁺ population detected after infection includes a contribution from ongoing microbiota-driven responses, it is largely infection-induced. Interestingly, YFP^+^ populations were spatially segregated within the gut: GC B cells were primarily localized to Peyer’s patches and mes LNs; PCs were mainly distributed throughout the small intestinal villi and large intestinal crypts; and MBCs were detected in both inductive and effector sites (Figures 1J-1N and S1G-S1H). In both the small and large intestine, PCs and MBCs were present at an approximately 80:20 ratio, resulting in a PC/MBC ratio significantly greater than 1 and revealing that the gut B cell memory compartment, in marked contrast to the lung, is largely dominated by PCs (Figures 1J-1M). Small and large intestinal PCs were antigen-specific, as they recognized *Salmonella* lipopolysaccharide (LPS) (Figure 1O). Notably, these PCs remained functionally active over extended periods, as LPS-specific antibody-secreting cells were detected in the lamina propria and LPS-specific antibodies were present in the feces up to 250 days post-infection (Figures S1I–S1J).

We next examined the functional implications of these divergent B cell memory strategies in lung and gut barrier tissues. To assess this, we primarily infected mice with either influenza PR8 or *Salmonella* SL3261 strains and rechallenged them at week 5 with influenza A virus HKx31 strain or *Salmonella* SL12023 strain. Four days after re-challenge, we measured the formation of antibody-secreting cells in the lungs, small intestine, and large intestine by ELISpot. Secondary infection led to a robust de novo formation of influenza-specific antibody-secreting cells in the lungs, whereas no significant increase in *Salmonella*-specific antibody-secreting cells was observed in the small and large intestines (Figures 1P and 1Q). These results are consistent with the notion that lung MBCs can rapidly differentiate into antibody-secreting cells to respond to influenza variants, whereas the gut appears to rely on antibodies produced by gut-resident PCs generated during the primary infection to cope with a secondary *Salmonella* encounter. While we cannot exclude the possibility that gut MBCs rapidly differentiate into PCs and replace pre-existing PCs following re-challenge, our overall results indicate that the lungs mount a rapid increase in antibody-secreting cells, whereas antibody-secreting cell numbers in the gut remain stable.

To determine whether the marked differences in B cell memory strategies observed in lung and gut barrier tissues are driven by the type of pathogen used or by intrinsic features of the barrier site, we infected Aid-EYFP mice with the same pathogen, *Klebsiella pneumoniae*, either intranasally or orally. Intranasal *Klebsiella* infection elicited an MBC-biased response in the lungs, whereas oral *Klebsiella* infection triggered a PC-biased response in the small and large intestines, resulting in PC/MBC ratios lower than 1 in the lungs and higher than 1 in the intestine (Figures 1R–1S). These findings show that the observed differences are not pathogen-related. Interestingly, analysis of *Klebsiella* LPS-specific PCs revealed higher numbers of LPS-specific antibody-secreting cells in the intestine than in the lungs (Figure 1T). Values were normalized to total cell numbers per organ to account for differences in organ size. Notably, the number of *Klebsiella*-specific PCs in the bone marrow was comparable between mice infected intranasally or orally, indicating that the reduced PC abundance in the lungs is not due to preferential migration of PCs to the bone marrow (Figures S1K–S1L). Instead, these results suggest that divergent B cell memory strategies are intrinsically programmed in the lungs and gut to cope with future pathogenic challenges.

### Selection in GCs shapes lung and gut B cell memory compartments

Next, we asked whether the contrasting B cell memory strategies observed in the lung and gut mucosa are associated with differential output of PCs and MBCs during GC selection. To address this, we tracked the abundance of pre-PCs and pre-MBCs in lung- and gut-associated tissues following intranasal or oral infection with *Klebsiella pneumoniae*. As pre-PCs (*18*) and pre-MBCs (*19*, *20*) represent transitional populations between the GC and the memory compartments, they can serve as proxies for GC selection outcomes. Pre-PCs were identified as CD19⁺IgD⁻GL7⁺Fas⁺CD38⁻CD39⁺ cells, whereas pre-MBCs were identified as CD19⁺IgD⁻GL7⁺Fas⁺CD38⁺CCR6⁺CXCR3⁺ cells (Figures S2A-S2F). Following infection, we observed a progressive accumulation of both pre-MBCs and pre-PCs in the lungs, med LNs, Peyer’s patches, and mes LNs (Figures 2A-2B and S2G-S2H). Notably, when we quantified the ratio of pre-PCs to pre-MBCs, this ratio was consistently higher at all time points in Peyer’s patches compared with the lungs, with a similar trend observed in mes LNs versus med LNs (Figures 2C and S2I). We further confirmed this pattern using the influenza and *Salmonella* infection models, in which the pre-PC/pre-MBC output ratio likewise remained elevated in gut tissues (Figures 2D–2F and S2J–S2L). Overall, these results suggest that the enrichment of PCs within the intestinal memory compartment is due, at least in part, to biased GC selection toward the pre-PC/PC lineage.

**Figure 2.**
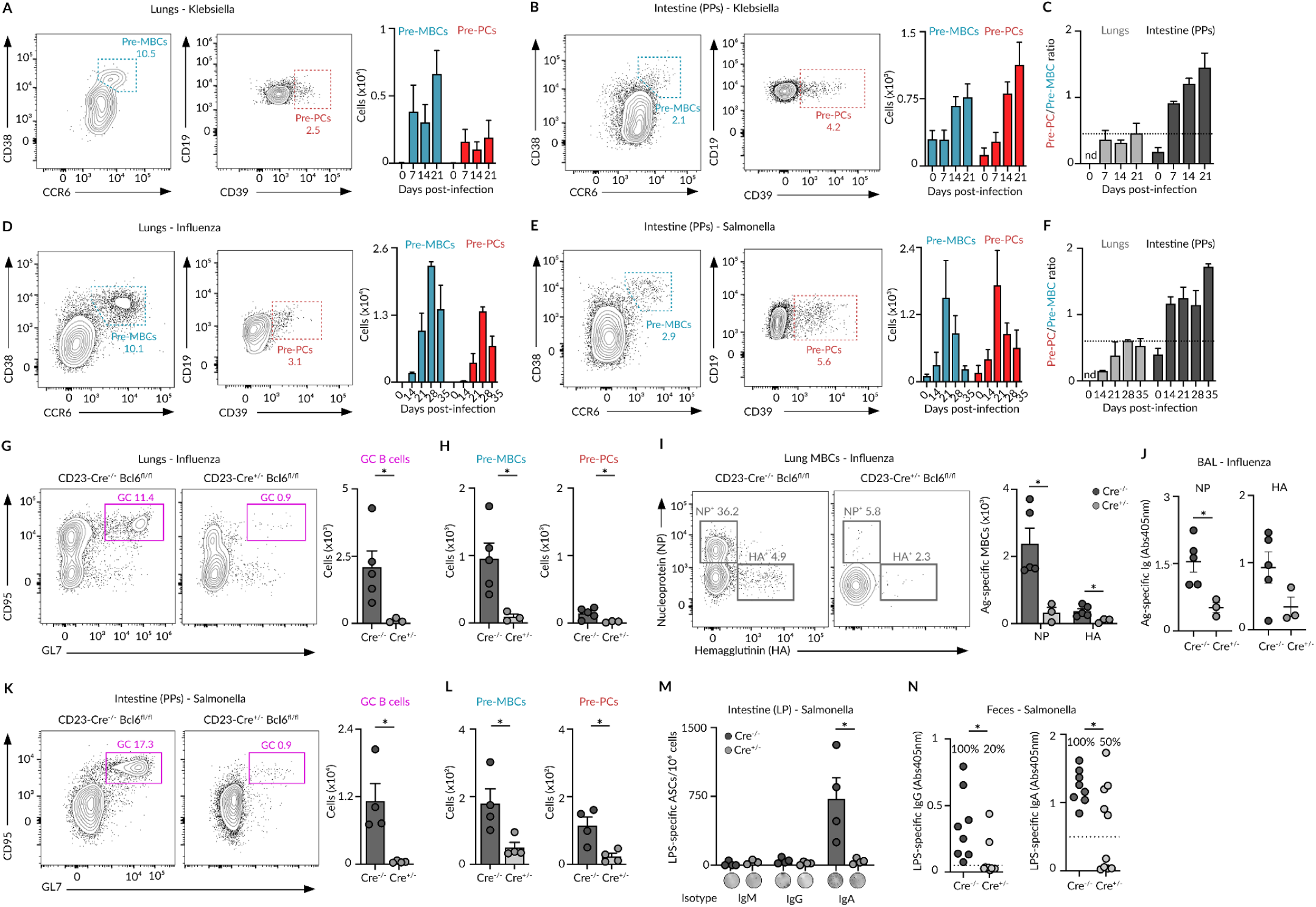
Differential selection in GC reactions shape mucosal memory compartments. (**A, B, D** and **E**) Flow cytometry plots and quantification of pre-MBCs (CD38^+^CCR6^+^) and pre-PCs (CD19^+^CD39^+^) within the GC reaction (CD95^+^GL7^+^) in (A and D) lungs and (B and E) intestine (Peyer’s patches) after (A and B) Klebsiella, (D) Influenza or (E) Salmonella infection. (**C** and **F**) Bar charts showing prePC:preMBC ratios in (C) lungs and (F) intestine after (C) Klebsiella or (F) Influenza/Salmonella infection. (**G** and **K**) Flow cytometry plots and quantification of GC B cells in (G) lungs and (K) intestine (Peyer’s patches) of CD23-cre Bcl6^fl/fl^ mice infected with influenza or Salmonella respectively. (**H** and **L**) Quantification of pre-MBCs (CD38^+^CCR6^+^) and pre-PCs (CD19^+^CD39^+^) within the GC reaction (CD95^+^GL7^+^) in (H) lungs and (L) intestine (Peyer’s patches) after Influenza or Salmonella infection. (**I**) Contour plots and quantification of lung MBCs binding to fluorescently-labeled NP and HA antigens. (**J**) Quantification of NP and HA-specific immunoglobulin levels in the bronchoalveolar fluid. (**M**) Enumeration of intestine LPS-specific IgM^+^, IgG^+^, and IgA^+^ antibody-secreting cells by ELISpot. (**N**) Quantification of LPS-specific immunoglobulin levels in fecal extracts. Percentage of responders is shown. All panels display quantification from one representative experiment out of three (mean ± s.e.m.). Each dot represents one mouse. t-test (**G**, **H**, **J**, **K**, **L** and **N**) and two-way ANOVA (**I** and **M**): *p<0.05.

During infection, B cell selection can occur not only within GCs but also through extrafollicular pathways (*21*). To assess the extent to which the observed differences in GC-associated memory commitment contribute to divergent memory strategies in the lungs and gut, we genetically disrupted GC formation by deleting the transcription factor Bcl6 in B cells (*22*). To this end, we crossed Bcl6^fl/fl^ mice with either CD19-Cre or CD23-Cre lines and infected the resulting animals with influenza virus or *Salmonella*. In CD19-Cre^+/-^ Bcl6^fl/fl^ mice, incomplete deletion of *Bcl6* did not prevent GC formation or memory responses, likely due to selective expansion of B cells that escaped deletion (Figures S3A-S3H). In contrast, CD23-Cre^+/-^ Bcl6^fl/fl^ mice exhibited efficient deletion of *Bcl6* in B cells, resulting in a marked reduction of GC B cells in the lungs and med LNs following influenza infection (Figures 2G and S3J). Disruption of GC formation led to a strong reduction in pre-PCs and pre-MBCs, indicating that these populations are GC-derived (Figures 2H and S3K). Consequently, this resulted in a reduction in lung-resident MBCs specific for influenza HA and NP, as well as decreased influenza-specific antibody levels in bronchoalveolar lavage fluid (Figures 2I-2J). Together, these data indicate that GC reactions are required for the generation of lung memory responses.

In the gut, CD23-Cre^+/-^ Bcl6^fl/fl^ mice also exhibited a strong reduction in GCs, pre-PCs, and pre-MBCs in Peyer’s patches and mes LNs following *Salmonella* infection (Figures 2K-2L and S3L-S3M). Despite the known capacity of *Salmonella* to elicit robust extrafollicular responses (*23*, *24*), we observed a significant reduction in LPS-specific gut PCs and fecal antibody titers in these mice (Figures 2M-2N), suggesting that GC output contributes substantially to the generation of gut-resident PCs even in the presence of active extrafollicular responses.

Altogether, these findings demonstrate that GC selection plays a central role in shaping mucosal memory compartments in both the lungs and gut.

### Mucosal memory selection highly relies on BCR isotype usage

Mucosal infections trigger GC reactions that not only comprise B cells with diverse BCR affinities, as observed in conventional immunization models, but also B cells expressing distinct BCR specificities and isotypes, including IgM, multiple IgG subclasses, and IgA (*25–28*). To assess the importance of these BCR features in mucosal memory selection, we took advantage of a series of mouse models targeting the mechanisms that generate BCR diversity. In these models, we tracked the formation of lung and intestinal PCs (CD95⁺CD39⁺CD98⁺) and bonafide MBCs (CD95⁺CD39^lo^CD38⁺CXCR3⁺) by flow cytometry (Figure S4) (*29*). We initially used AID-deficient mice, which generate a diverse BCR repertoire but cannot undergo isotype class switching or affinity maturation, although polyclonal populations may still exhibit differences in baseline affinity (Figures S5A-S5B). We found that AID deficiency led to a marked reduction in pre-PC output in the lungs following influenza infection, resulting in a lower pre-PC/pre-MBC ratio (Figures 3A and S5C). This impairment was accompanied by a strong reduction in the PC/MBC ratio in the lungs, as well as diminished generation of influenza-specific PCs (Figures 3B and S5D-S5E). Following *Salmonella* infection, we likewise observed a pronounced reduction in the pre-PC/pre-MBC ratio in Peyer’s patches, resulting in decreased PC/MBC ratios in the intestine and reduced formation of *Salmonella* LPS-specific PCs (Figures 3C-3D and S5F-S5I). These results indicate that affinity maturation and/or class-switch recombination are critical for B cell selection into the PC compartment during mucosal infection. To disentangle the contribution of these two processes, we used Aid-*G23S* mutant mice, which carry a point mutation in AID that selectively impairs somatic hypermutation while preserving class switching (Figures S6A-S6B) (*30*). Notably, pre-PC/pre-MBC ratios, PC/MBC ratios, and the generation of pathogen-specific PCs were comparable between wild-type and Aid-*G23S* mice following influenza or *Salmonella* infection (Figures 3E–3H and S6C-S6I). These results indicate that limiting affinity maturation alone does not significantly impact mucosal B cell selection into the PC compartment in the models tested.

**Figure 3.**
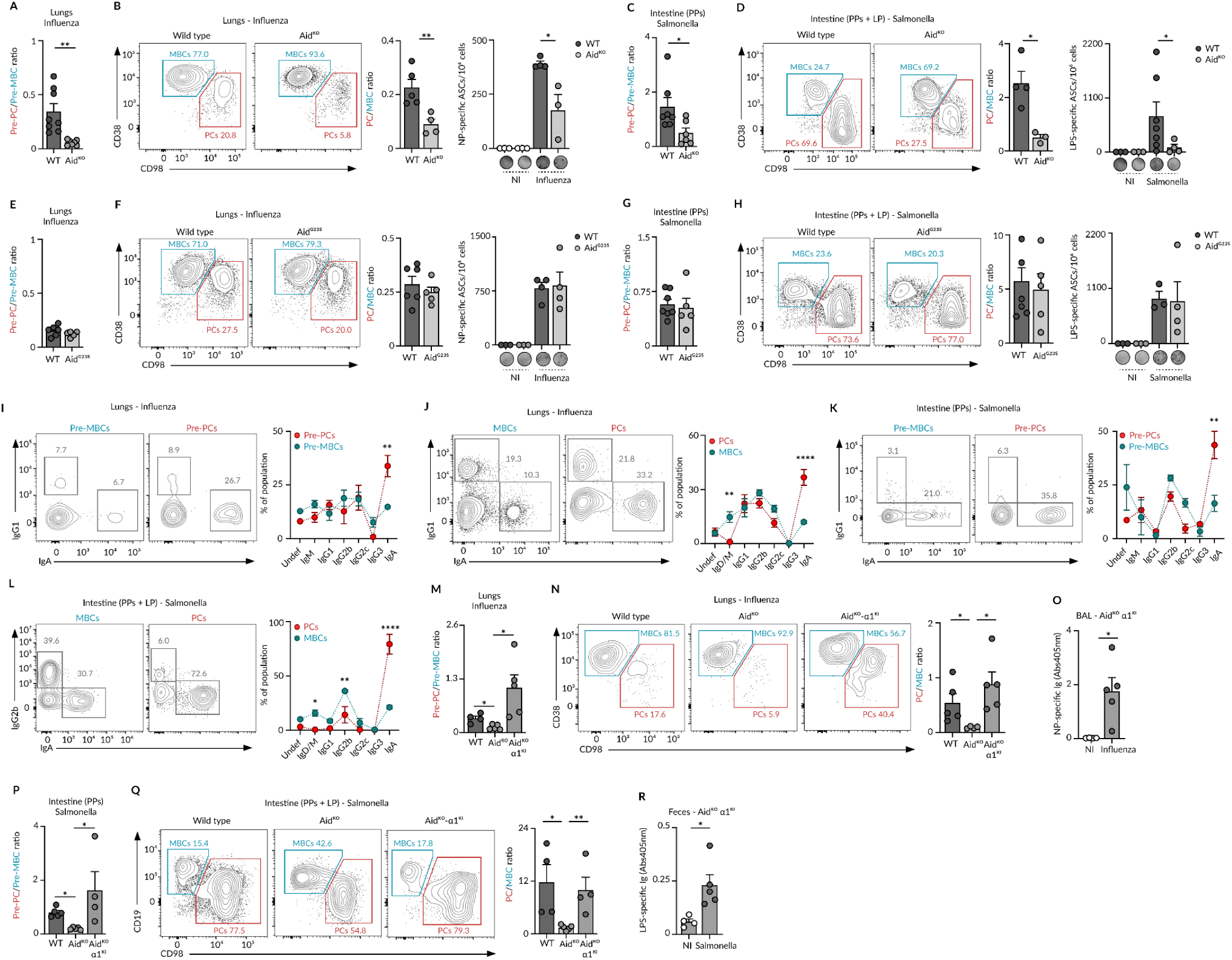
Mucosal memory selection relies on IgA BCR expression rather than affinity. (**A** and **C**) Bar charts showing pre-PC:pre-MBC ratios in lungs or intestine of influenza- or Salmonella-infected wild type and *Aid*-KO mice. (**B** and **D**) Flow cytometry plots displaying MBC (CD38^+^CD98^-^) and PC (CD38^low^CD98^+^) distribution in the lungs or intestine of influenza or Salmonella-infected wild type and *Aid*-KO mice. Bar charts show PC:MBC ratio and the enumeration of lung NP- or gut LPS-specific antibody secreting cells by ELISpot. (**E** and **G**) Bar charts showing pre-PC:pre-MBC ratios in lungs or intestine of influenza- or Salmonella-infected wild type and *Aid*(G23S) mice. (**F** and **H**) Flow cytometry plots displaying MBC (CD38^+^CD98^-^) and PC (CD38^low^CD98^+^) distribution in the lungs or intestine of influenza- or Salmonella-infected wild type and *Aid*(G23S) mice. Bar charts show PC:MBC ratio and the enumeration of lung NP- or gut LPS-specific antibody secreting cells by ELISpot. (**I** and **K**) Contour plots and quantification of isotype distribution within pre-MBCs and pre-PCs in (I) lungs of influenza-infected mice and (K) intestine of *Salmonella*-infected mice. (**J** and **L**) Contour plots and quantification of isotype distribution within MBCs and PCs in (J) lungs of influenza-infected mice and (L) intestine of *Salmonella*-infected mice. (**M** and **P**) Bar charts showing pre-PC:pre-MBC ratios in (M) lungs or (P) intestine of influenza- or Salmonella-infected wild type, *Aid*-KO and *Aid*-KO-ɑ1KI mice. (**N** and **Q**) Flow cytometry plots displaying MBC versus PC distribution in (N) lungs of influenza-infected and (Q) intestine of *Salmonella*-infected wild type, *Aid*-KO and *Aid*-KO-ɑ1KI mice. Bar charts show PC:MBC ratios. (**O**) Quantification of NP-specific immunoglobulin levels in the bronchoalveolar fluid of influenza-infected *Aid*-KO-ɑ1KI mice. (**R**) Quantification of LPS-specific immunoglobulin levels in fecal extracts of Salmonella-infected *Aid*-KO-ɑ1KI mice. All panels display quantification from one representative experiment out of three (mean ± s.e.m.). Each dot represents one mouse. t-test (**A**-**H, O** and **R**), one-way ANOVA (**M**, **N, P** and **Q**), two-way ANOVA (**I-L**): *p<0.05, **p<0.01, and ****p<0.0001.

Based on these results, we hypothesized that class switching toward distinct BCR isotypes may play a dominant role in directing B cell memory selection at mucosal sites. To test this, we measured the BCR isotype distribution in pre-PCs and pre-MBCs in lungs and med LNs after influenza infection. We found that IgG⁺ cells contributed to pre-MBC and pre-PC compartments to a similar extent, whereas B cells that had class-switched to IgA were highly enriched in the pre-PC population (Figures 3I and S7A-S7B). This bias toward pre-PC differentiation among IgA⁺ cells was reflected by their preferential accumulation in the PC compartment (Figures 3J and S7C-S7D). We observed a similar enrichment of IgA^+^ cells in the pre-PC and PC compartments in the intestine following *Salmonella* infection (Figures 3K-3L and S7E-S7G). These results suggest that the IgA BCR favors B cell selection into the PC program. To directly test this idea, we used α1KI transgenic mice, in which IgA expression is genetically driven by replacement of the IgM constant region with the IgA constant region (*31*). We further crossed α1KI mice with Aid-KO animals to block somatic hypermutation and class switching to other isotypes (Figures S8A-S8B). We infected wild-type, Aid-KO, and Aid-KO/α1KI mice with either influenza virus or *Salmonella* and measured B cell selection into the pre-PC and pre-MBC compartments, as well as memory populations. We found that forced expression of the IgA BCR partially restored the impaired pre-PC selection and PC differentiation observed in Aid-KO mice in both lungs and gut (Figures 3M–3R and S8C–S8I). Together, these results support the notion that the IgA BCR favors B cell differentiation into PCs independently of affinity maturation and indicate that mucosal memory selection is strongly influenced by BCR isotype usage.

### Mucosal memory selection integrates microbial cues via the IgA BCR

Given the preferential differentiation of IgA^+^ B cells into the PC compartment, we asked whether the PC-biased response in the gut reflects increased usage of the IgA BCR in gut-associated GCs compared with lung GCs. To test this, we analyzed IgA isotype usage among GC B cells in the lungs and intestine following intranasal or oral *Klebsiella* infection. We found that class switching to IgA was approximately tenfold more frequent in Peyer’s patches than in the lungs (Figures 4A and S9A). A similar enrichment of IgA usage in gut versus lung GC responses was also observed in the influenza and Salmonella infection models (Figures 4B and S9B), suggesting that elevated IgA class switching in gut-associated GCs contributes to enhanced PC output in this tissue. To directly test the impact of the IgA BCR in PC differentiation, we infected wild-type and Igha-deficient mice with influenza virus or Salmonella and analyzed pre-PC/pre-MBC output and memory responses. Igha-deficient mice showed a significant reduction in the pre-PC/pre-MBC ratio, accompanied by decreased total PC output and fewer pathogen-specific PCs in both infection models (Figures 4C-4F and S9C-S9I). These mice also exhibited increased numbers of IgM-secreting cells, consistent with a compensatory response in the absence of IgA (Figures S9J-S9K). Importantly, a similar reduction in PC commitment was observed after both intranasal and oral *Klebsiella* infection across lung and intestinal compartments (Figures S9L-S9O). Altogether, these findings support a model in which the IgA BCR favours PC differentiation and in which the extent of IgA usage during GC reactions shapes mucosal memory output during infection.

**Figure 4.**
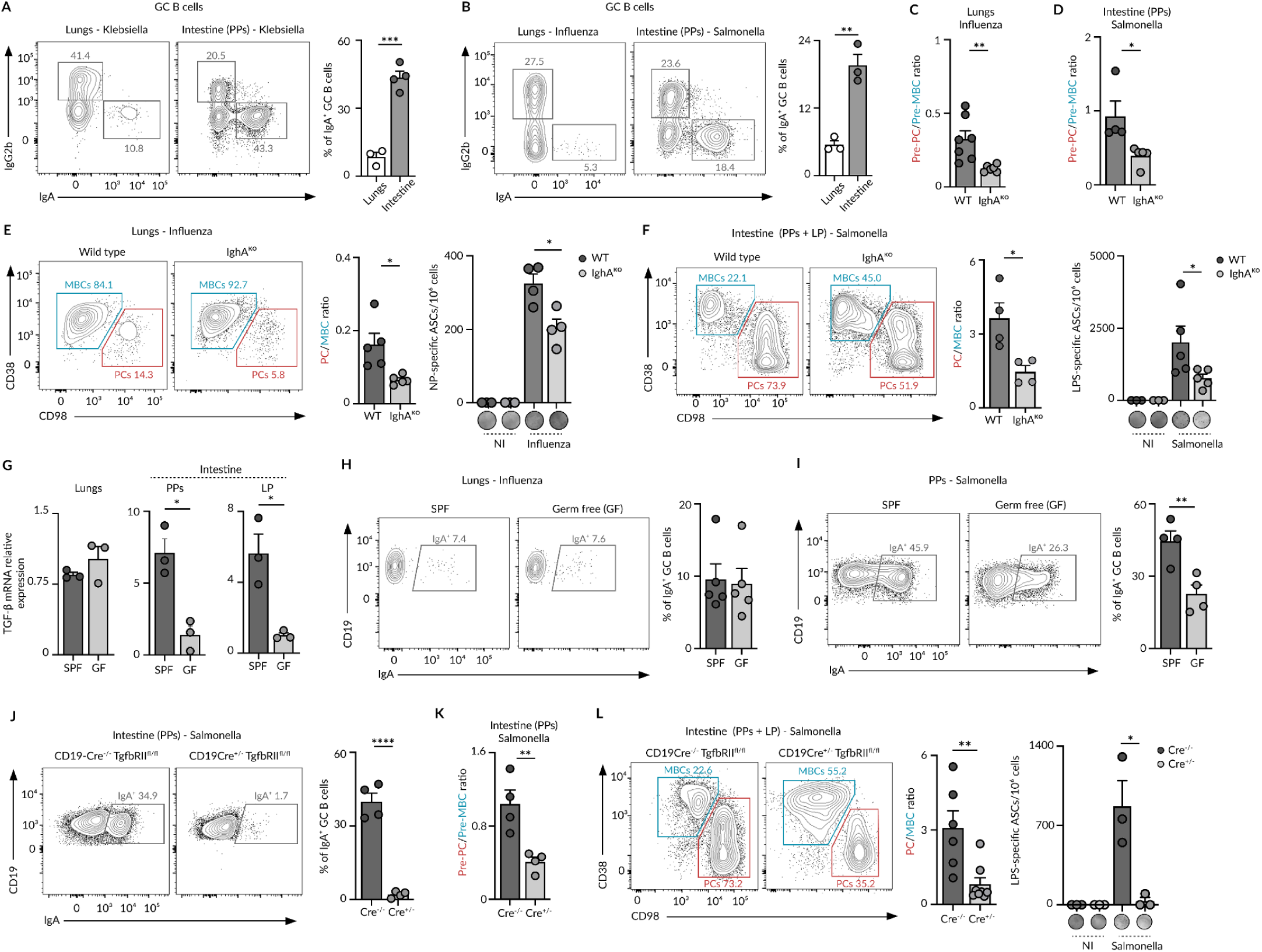
The intestinal microbiota favors the TGFβ-IgA-PC axis. (**A** and **B**) Flow cytometry plots and quantification of IgA BCR isotype usage within the GC population in lungs or intestine (Peyer’s patches) after (A) *Klebsiella* or (B) Influenza/*Salmonella* infection (day 14). (**C** and **D**) Bar charts show prePC:preMBC ratios in (C) lungs and (D) intestine (Peyer’s patches) of wild type and *IghA*-KO mice infected with influenza virus or *Salmonella*, respectively. (**E** and **F**) Flow cytometry plots displaying MBC (CD38^+^CD98^-^) and PC (CD38^low^CD98^+^) distribution in (E) lungs or (F) gut of influenza- or Salmonella-infected wild type and *IghA*-KO mice. Bar charts show PC:MBC ratio and the enumeration of lung NP- or intestinal LPS-specific antibody secreting cells by ELISpot. (**G**) Quantification of TGFβ mRNA levels in lung and intestine of SPF and Germ free mice by qPCR. Cytokine expression was normalized to β-actin. (**H** and **I**) Contour plots and quantification of IgA expression by GC B cells in (H) lungs and (I) intestine of influenza or Salmonella-infected mice. (**J**) Flow cytometry plots and quantification displaying IgA expression by GC B cells in the intestine (Peyer’s patches) of *Salmonella*-infected CD19-cre TgfbRII^fl/fl^ mice. (**K**) Bar chart shows pre-PC:pre-MBC ratios in the intestine of Salmonella-infected CD19Cre- TgfbRII^fl/fl^ mice. (**L**) Contour plots and quantification of intestinal MBCs and PCs in the intestine of CD19Cre- TgfbRII^fl/fl^ mice infected with *Salmonella*. Bar charts represent PC:MBC ratios and the enumeration of intestinal LPS-specific antibody secreting cells by ELISpot. All panels display quantification from one representative experiment out of three (mean ± s.e.m.). Each dot represents one mouse. t-test (**A**-**L**): *p<0.05, **p<0.01, ***p<0.001 and ****p<0.0001.

Class switching toward IgA is primarily driven by TGF-β, a cytokine produced in response to microbial colonization (*32–36*). We hypothesized that the high microbiota biomass in the intestine contributes to the elevated IgA class switching observed in gut-associated GC reactions. To test this, we compared TGF-β levels in germ-free and specific-pathogen-free (SPF) mice. While TGF-β levels in lungs were similar between the two groups, Peyer’s patches and lamina propria of germ-free mice showed significantly reduced TGF-β levels compared to SPF controls (Figure 4G). Interestingly, when we infected germ-free and SPF mice with influenza virus or Salmonella, IgA class switching in the lungs occurred at comparable levels in both groups (Figure 4H). In contrast, IgA class switching in Peyer’s patches was markedly reduced in germ-free mice (Figure 4I). These findings suggest that the elevated microbial load in the gut promotes TGF-β production and IgA class switching. To directly test whether the microbiota–TGF-β–IgA axis favors the PC-dominated gut memory response through B cell-intrinsic TGF-β sensing, we generated mice lacking TGF-β receptor II specifically in B cells (CD19-Cre^+/-^ Tgfbr2^fl/fl^). We then infected CD19-Cre^+/-^ and Cre^-/-^Tgfbr2^fl/fl^ mice with Salmonella. In CD19-Cre^+/-^ Tgfbr2^fl/fl^ mice, IgA^+^ GC B cells were absent in Peyer’s patches, consistent with the requirement for TGF-β signaling in IgA class switching (Figure 4J). This was accompanied by a reduced pre-PC/pre-MBC ratio (Figures 4K and S10A). Notably, the strong PC bias normally observed in the gut was partially reversed when B cells lacked TGF-β signaling, even in the presence of microbiota (Figures 4L and S10B-S10D). Furthermore, the number of Salmonella LPS–specific PCs was reduced in the absence of TGF-β signaling in B cells (Figure 4L). Together, these findings support a model in which high microbial biomass in the gut enhances TGF-β production and IgA class switching, thereby favouring PC differentiation. Overall, B cell selection in mucosal tissues integrates site-specific microbial cues through TGF-β signaling and IgA BCR usage, shaping the memory outcome upon infection.

### The IgA BCR relies on its cytosolic tail to amplify BCR signalling

The strength of BCR signaling plays a pivotal role in determining B cell fate upon antigen recognition (*37–39*). Recent studies further demonstrated that the IgA BCR transmits signals of different magnitude compared to other isotypes (*40*, *41*). Based on this, we hypothesized that selectively altering IgA BCR signaling could modulate mucosal B cell selection and the PC:MBC output during infection. To test this, we aimed to selectively manipulate IgA BCR signaling without affecting other isotypes. Class-switched BCRs (IgG and IgA), unlike those on naïve B cells (IgD and IgM), contain a conserved cytoplasmic tail across species (*42*). While previous studies have shown that the IgG tail enhances BCR signaling (*43*, *44*), the functional role of the IgA tail, and whether it contributes to signal strength, remains unknown. To begin addressing this, we genetically engineered GC-like IgG2a^+^ A20 B cells to express an IgA BCR by introducing CRISPR-Cas9 guide RNAs targeting the Sγ2a-3’ and Sα-3’ switch regions (*45*) (Figure 5A). This strategy was highly efficient in forcing class-switching from IgG to IgA while preserving surface BCR expression, as confirmed by anti-kappa light chain staining (Figures 5B-5C). We then generated IgA^+^ B cells lacking the cytoplasmic tail sequence (RGPFGSKEVPQY) using CRISPR-Cas9. To facilitate identification of successful recombinants, we inserted P2A and CD90.1 coding sequences into both wild-type and Δtail constructs (Figure 5D). Notably, deletion of the tail did not affect membrane expression of IgA (Figures 5E–5F).

**Figure 5.**
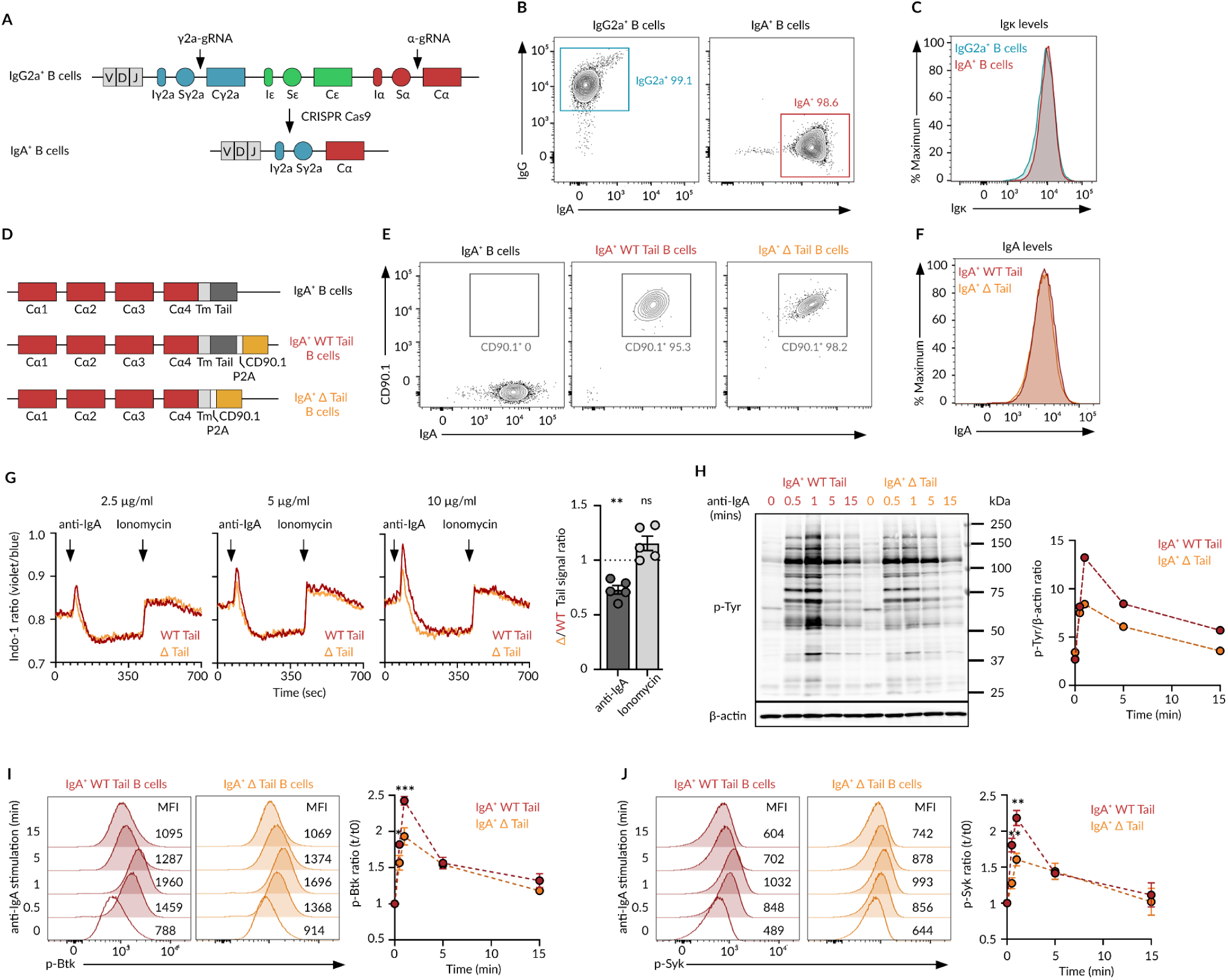
The IgA BCR requires its cytosolic tail for efficient signal transduction. (**A**) Schematic representation of the strategy used to force class-switching of A20 B cells from IgG2a to IgA with CRISPR/Cas9. (**B**) Contour plots show IgG2a versus IgA expression 21 days after transfection and clonal dilution of cells engineered as in (A). (**C**) Histogram shows Κappa light chain levels in IgG2a^+^ and IgA^+^ A20 B cells. (**D**) Schematic strategy used to remove the cytosolic tail from IgA^+^ A20 B cells. (**E**) Contour plots show the expression of IgA BCR and CD90.1 in WT and Δtail A20 B cells generated in (D). (**F**) Histogram shows IgA BCR expression at the surface of IgA WT and Δtail A20 B cells. (**G**) Ca^2+^ mobilization in IgA^+^ WT (red) and IgA^+^ Δtail (orange) A20 B cells stimulated with increasing concentrations of anti-mouse IgA and ionomycin at indicated time points. Quantifications were done by dividing the maximum value of the peak of the IgA ΔTail cells stimulation by the maximum value of the peak of the IgA WT Tail cells, after either anti-IgA or Ionomycin stimulation. Each dot represents one experiment. (**H**) Immunoblot analysis and quantification of lysates from IgA^+^ WT (red) and IgA^+^ Δtail (orange) A20 B cells, either untreated or stimulated for various times with anti-IgA. Membranes were probed with anti-phosphotyrosine (top) and anti-β-actin (bottom). Quantification of one representative experiment out of three is shown. (**I** and **J**) Phospho-flow cytometry histograms and quantification of (I) Btk and (J) Syk MFI in IgA^+^ WT (red) and IgA^+^ Δtail (orange) B cells after stimulation with anti-mouse IgA. MFI values were relativized to time=0. Quantifications of three experiments are shown (mean ± s.e.m.). For panel G, we perform a t-test to calculate whether the Δ/WT tail ratio was different to 1, Two-way ANOVA (I and J): *p<0.05, **p<0.01 and ***p<0.001

To evaluate the functional relevance of the IgA tail in BCR signaling, we performed a series of assays. First, calcium flux analysis revealed that Δtail IgA^+^ B cells displayed reduced calcium mobilization compared to wild-type cells following cross-linking with increasing concentrations of anti-IgA antibody, while both B cells responded similarly to ionomycin (Figure 5G). Next, we assessed global phosphorylation using anti-phosphotyrosine (4G10) western blotting and observed a marked reduction in the phosphorylation landscape of Δtail IgA^+^ B cells upon BCR cross-linking (Figure 5H). Finally, phospho-flow cytometry showed significantly reduced phosphorylation of early signaling molecules Btk and Syk in Δtail IgA^+^ B cells upon stimulation (Figures 5I-5J). Altogether, these findings demonstrate that the cytoplasmic tail of the IgA BCR is required for effective signaling upon antigen engagement. Its removal selectively impairs IgA BCR signaling, providing a tool to specifically modulate this pathway without affecting other isotypes.

### The IgA BCR cytosolic tail shapes mucosal B cell memory responses

To assess to which extent IgA BCR signalling can impact B cell memory selection, we generated new transgenic mice in which we introduced a P2A-tdTomato cassette at the end of the IghA coding sequence to track IgA^+^ cells (Igha-tdTom) and subsequently deleted the cytosolic tail domain (Igha-Δtail-tdTom) (Figure 6A). Labeling was highly efficient in both mouse strains, as IgA^+^ GCs, MBCs and PCs expressed high levels of tdTomato in lungs and gut upon infection (Figures S11A and S11B). Yet, some IgA^-^ cells, in particular PCs, expressed considerable levels of tdTomato, although to a lesser extent than IgA^+^ cells (Figures S11A and S11B). This phenomenon might be due to expression of IgA germline transcripts in IgG^+^ cells and/or switch transcripts from the nonproductive heavy-chain allele, as previously observed in Cγ1-Cre, IgE-GFP mice and IgE-tdTomato mice (*46–48*).

**Figure 6.**
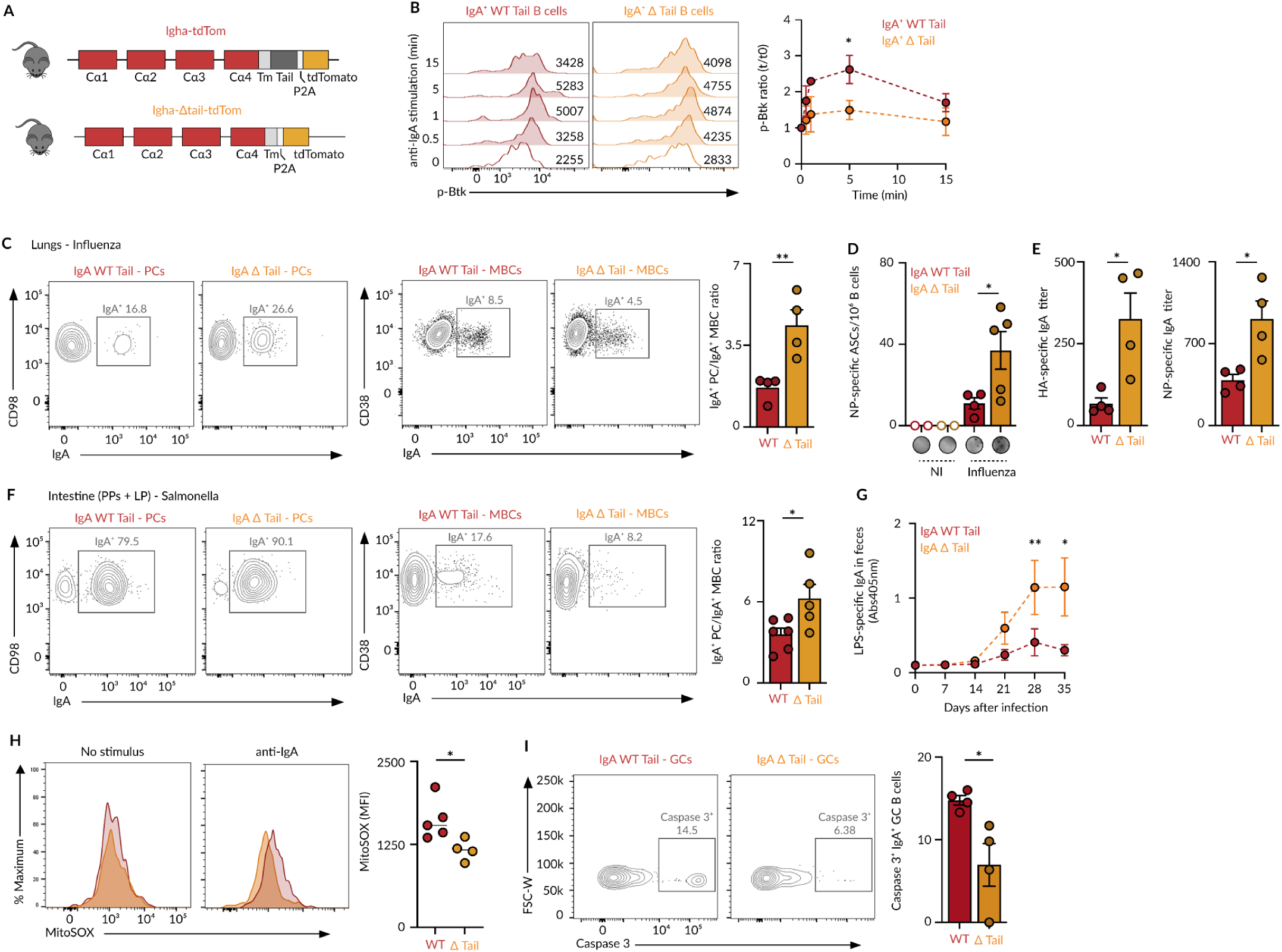
The IgA cytosolic tail balances PC entry by modulating BCR signaling. (**A**) Schematic representation of Igha-tdTom and Igha-Δtail-tdTom mouse lines. (**B**) Phospho-flow cytometry histograms and quantification of Btk MFI in IgA^+^ WT (red) and IgA^+^ Δtail (orange) B cells from Peyer’s patches after stimulation with anti-mouse IgA. MFI values were relativized to time=0. Quantification of three experiments is shown (mean ± s.e.m.). (**C** and **F**) Contour plots displaying IgA^+^ cells within MBC and PC populations in (**C**) lungs and (**F**) intestine of Igha-tdTom and Igha-Δtail-tdTom mice infected with influenza virus or *Salmonella*, respectively. Quantifications show IgA^+^ PC: IgA^+^ MBC ratios. (**D** and **E**) Quantification of (D) NP-specific PCs in the lungs and (E) HA- and NP-specific IgA levels after influenza infection. (**G**) Quantification of LPS-specific IgA antibodies in the feces after Salmonella infection. Quantification of two experiments is shown (mean ± s.e.m.) **(H)** Flow cytometry histograms and quantification of MitoSOX fluorescence in IgA^+^ WT (red) and IgA^+^ Δtail (orange) GC B cells incubated with or without anti-IgA. **(I)** Flow cytometry analysis and quantification of Caspase-3 staining in IgA^+^ WT (red) and IgA^+^ Δtail (orange) GC B cells. Unless stated, panels display quantification from one representative experiment out of three (mean ± s.e.m.). Each dot represents one mouse. t test (C, D, E, F, H and I) and 2-way ANOVA (B and G): *p<0.05 and **p<0.01.

We next tested whether removal of the IgA cytoplasmic tail affects BCR signaling in primary B cells. To do this, we stimulated IgA^+^ GC B cells from Igha-tdTom and Igha-Δtail-tdTom Peyer’s patches with anti-IgA antibody and measured Btk phosphorylation by phospho-flow cytometry. In line with our earlier results from GC-like B cell lines, deletion of the IgA cytoplasmic tail significantly reduced Btk phosphorylation upon BCR stimulation, confirming that the IgA BCR requires its cytosolic tail to effectively transduce signaling (Figure 6B). We then assessed the functional consequences of reduced BCR signaling on B cell selection. To address this, we infected Igha-tdTom and Igha-Δtail-tdTom mice with influenza virus or Salmonella and analyzed B cell commitment to the pre-MBC and pre-PC pathways, as well as memory responses. Surprisingly, IgA⁺ cells from Igha-Δtail-tdTom mice were more efficiently selected into the pre-PC pathway than those from Igha-tdTom counterparts in both the lungs and gut (Figures S11C-S11D). This led to an enhanced accumulation of lung and intestinal PCs, significantly increased numbers of pathogen-specific PCs and higher IgA levels in serum and feces (Figures 6C-6G and S11E-S11I).

To further validate our findings in a competitive setting, we took advantage of allelic exclusion, a mechanism by which B cells express only one heavy chain locus during development. We analyzed heterozygous Igha-tdTom^+^/^−^ and Igha-Δtail-tdTom^+^/^−^ mice, which generate two genetically distinct populations of IgA^+^ B cells: one expressing wild-type IgA and the other expressing either IgA-tdTom or IgA-Δtail-tdTom. Using tdTomato fluorescence, we were able to distinguish IgA^+^ cells derived from the modified versus wild-type alleles (Figures S12A-S12B). We infected these heterozygous mice with influenza or Salmonella and analyzed IgA^+^ cells in the memory compartments 35 days later. In both lungs and gut, we observed that Igha-Δtail-tdTom^+^ cells were consistently enriched in the PC compartment compared to Igha-tdTom^+^ cells (Figures S12A-S12B). These results exclude the possibility that external factors are responsible for the enhanced PC output of IgA^+^ B cells lacking the cytosolic tail. Instead, our results support the notion that intrinsic differences in BCR signaling underlie divergent memory fates.

Although these results were initially unexpected, recent studies have shown that BCR signaling in GC B cells must be tightly regulated to prevent excessive accumulation of reactive oxygen species (ROS) and subsequent cell death (*49*). For instance, enhanced BCR signaling resulting from phosphatase deletion in IgG^+^ GC B cells leads to increased oxidative stress, which impairs PC differentiation (*49*, *50*). To determine whether the increased PC output observed in Igha-Δtail mice was linked to reduced oxidative stress, we measured mitochondrial ROS levels using MitoSOX staining and apoptosis using caspase-3 staining in IgA^+^ GC B cells of Igha-tdTom and Igha-Δtail-tdTom mice. We found that IgA^+^ cells lacking the cytoplasmic tail accumulated less ROS upon BCR cross-linking and exhibited fewer caspase-3^+^ cells, indicating reduced apoptosis (Figures 6H-6I). Altogether, our findings suggest that the cytoplasmic tail of the IgA BCR modulates signaling strength and fine-tunes B cell selection during mucosal infection.

## Discussion

B cell memory responses at barrier tissues provide rapid and efficient protection against mucosal pathogens (*51–55*). In our study, we uncovered that distinct B cell memory strategies evolved in lungs and gut to counteract recurrent pathogens. Memory selection is skewed towards MBCs in the lungs while a strong PC-bias exists in the gut. We found that this divergence is linked to differential BCR isotype usage across barrier tissues rather than to affinity. In the gut, the high microbiota content fosters a regulatory environment rich in TGF-β and Treg cells that prevent excessive immune reactions to commensal bacteria while promoting B cell class switching towards IgA (*33*, *56*). In turn, the IgA BCR favors B cell entry into the PC program by fine-tuning downstream signaling. These results support a model in which mucosal memory selection integrates tissue-specific microbial cues by highly relying on BCR isotype usage rather than affinity. Furthermore, our findings raise the possibility that the IgA- and PC-biased memory response in the gut is a by-product of evolutionary pressures that have shaped a regulatory environment in the gut to limit the immune system’s activation against the resident microbiota.

What features of the IgA BCR favour B cell differentiation into PCs remains an important open question. Previous studies have reported differences in the relative magnitude of IgA-mediated signaling compared with IgG, with one study observing higher and another lower signaling outputs downstream of the IgA BCR (*40*, *41*). The reasons underlying these observations are not yet fully understood and may reflect differences in experimental context, including the anatomical origin of the analyzed B cells (intestinal versus circulating) as well as the use of distinct anti-BCR antibodies to trigger signaling. Future studies comparing B cells of matched specificity, affinity, and tissue origin stimulated with cognate antigen will help refine our understanding of how IgA BCR signaling compares to other isotypes.

We initially hypothesized that dampening IgA BCR signaling would impair PC differentiation. Unexpectedly, we observed the opposite outcome: genetic deletion of the IgA cytoplasmic tail reduced BCR signaling yet enhanced commitment to the PC fate. Although initially counterintuitive, this finding is consistent with recent studies demonstrating that excessive BCR signaling in GC B cells can be detrimental to their selection and PC differentiation due to increased ROS production and elevated intracellular calcium levels (*49*, *50*, *57*, *58*). In these models, BCR-induced ROS accumulation can be partially counterbalanced by T cell help; however, when signaling exceeds a certain threshold, T cell-derived survival signals become insufficient, resulting in apoptosis (*49*). In this context, we propose that removal of the IgA cytoplasmic tail attenuates BCR signaling strength, thereby limiting ROS accumulation and promoting B cell survival even under conditions of limited T cell help. Such a mechanism would favor PC differentiation but may do so at the expense of stringent affinity-based selection. Accordingly, IgA⁺ PCs generated in the absence of the cytoplasmic tail may display reduced affinity compared with their wild-type counterparts. This possibility will be directly addressed in future studies using single-cell BCR sequencing combined with affinity measurements to determine how reduced IgA BCR signaling impacts the quality of the PC compartment. Beyond the cytoplasmic tail, additional molecular features of the IgA BCR may also shape B cell responsiveness and fate decisions. For example, the nanoscale organization of the BCR within the plasma membrane differs among IgM, IgD, and IgG isotypes (*59*, *60*), but remains unexplored for IgA. Moreover, the unique structural properties of the IgA hinge region may further contribute to its functional specialization. These mechanisms are currently under investigation in our laboratory and represent important directions for future research.

Recent studies have advanced our understanding of how homeostatic and infection-induced IgA^+^ PCs are generated in the gut mucosa. At steady state, gut GCs continuously produce new IgA^+^ PCs that seed the lamina propria and differentiate into both short- and long-lived populations, with clonal renewal rather than long-term cellular persistence shaping intestinal IgA responses (*61*). Additional studies have highlighted the complexity of IgA^+^ PC responses during challenge. For example, repeated mucosal immunization with ovalbumin and cholera toxin can drive sequential class switching, with a fraction of IgA^+^ B cells arising from IgG1^+^ precursors (*62*). Moreover, intestinal viral infection induces interferon-γ dependent CXCR3 expression on IgA^+^ PCs, promoting their migration to the lamina propria (*63*). Our study provides insight into why the gut memory compartment is enriched in IgA^+^ PCs compared with other barrier tissues such as the lungs, and further shows that attenuating IgA BCR signaling through deletion of the cytoplasmic tail enhances PC differentiation. Importantly, although our data support a role for the IgA BCR in biasing GC output toward the PC fate, IgA expression alone is not sufficient to dictate memory outcomes across tissues. Indeed, IgA-deficient mice still display a predominance of PCs in the gut, and IgA-overexpressing mice retain a relative enrichment of MBCs in the lung, indicating that additional factors contribute to shaping B cell fate decisions. We therefore propose that IgA usage favors PC commitment compared to other isotypes, but acts within a broader, multifactorial network of local environmental cues and signaling pathways that collectively determine MBC versus PC differentiation in mucosal tissues.

In our study, we consider MBCs and PCs as final separate entities. However, previous work has shown that the PC pool can receive contributions from MBCs in the context of responses to microbiota or dietary antigens, suggesting that MBC and PC fates can be interdependent (*16*, *64*). In such settings of persistent antigen exposure, memory B cells can re-enter differentiation pathways and contribute to the PC compartment. In contrast, during acute infections where the pathogen is cleared, reactivation of MBCs and their differentiation into PCs may be less frequent. Although our data do not exclude a contribution of MBC-derived PCs, the predominant differences we observe at the GC selection level, particularly in pre-PC and pre-MBC populations, are more consistent with altered GC output rather than secondary MBC reactivation.

Finally, our results prompt us to consider the potential benefit of having two alternative memory strategies in different barrier tissues. In the gut, where we are occasionally exposed to pathogens through poisoned food and water, including *E. coli*, *Salmonella*, *Shigella*, *Rotavirus* and *Norovirus*, a memory response based on antibodies produced by lamina propria-resident PCs might be beneficial for the rapid opsonization and elimination of these pathogens from the gut lumen upon re-encounter (*13*, *65*). Conversely, in the lungs, we are frequently exposed to airborne viruses that rapidly spread through the community and generate new variants every season. In this context, it is advantageous to have a population of tissue-resident MBCs with broader antigen affinities that can recognize viral variants, thereby complementing the humoral immunity provided by lung PCs (*29*, *66*). These different B cell memory strategies employed by the lungs and the gut are key factors to consider when developing effective nasal and oral vaccines against mucosal pathogens.

## Acknowledgements

We thank Rejane Rua, Achille Broggi and Romain Roncagalli for analysis and interpretation of confocal images and signaling experiments. We thank Myriam Baratin, Pierre Golstein and Philippe Naquet for scientific discussions and feedback on the manuscript. We thank Manon Fabregue, Solene Mathieu and Capucine Guerry for technical support. We thank Bernard Malissen and CIPHE-GEMTIS Consortium investigators for the development of Igha-tdTom and Igha-Δtail-tdTom mice. We thank Claude-Agnès Reynaud and Jean Claude Weill for providing Aicda-CreERT2 mice. We thank Tasuku Honjo and Daniel Pinschewer for providing *Aid*(G23S) mice. We thank Julien Marie for providing Tgfbr2^fl/fl^mice. We thank Ronan Le Goffic for providing influenza virus strains. We thank David Holden for providing *Salmonella* sp. strains. We thank Régis Tournebize for providing *Klebsiella* sp. strains. We thank the flow cytometry and imaging (Imagimm) platforms for technical support and the biological resource unit for the breeding of animals (CIML).

## Author contributions

MG conceived and supervised the study. MPH, SO and MG designed the experiments. MPH, SO, LB, MM, AT, AZ, CG and MG performed the experiments. FF developed Igha-tdTom and Igha-Δtail-tdTom mice. ES, ML and MC provided materials. ES, ML, MC and CG provided discussion and edited the paper. MG, MPH and SO analyzed the data, made the figures and wrote the paper.

## Data and materials availability

All data are available in the main text or the supplementary materials.

**Figure S1.**
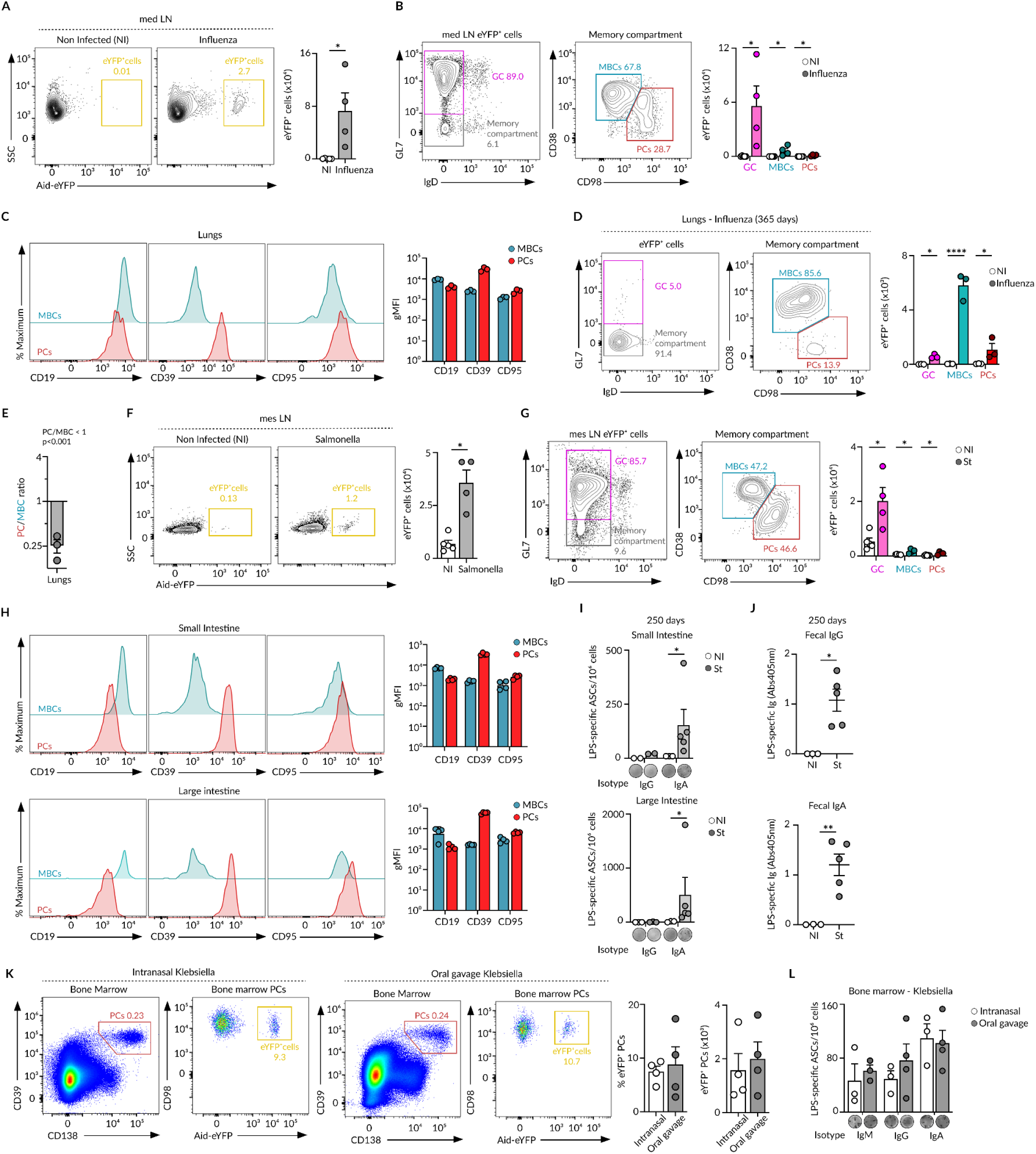
Characterization of mucosal B cell memory compartments. (**A**) Flow cytometry plots of med LN cells in non-infected (NI) or influenza-infected AID-EYFP mice and treated as in Figure 1A (**B**) Flow cytometry plots display the gating strategy of YFP^+^ GC B cells (GL7^+^), memory cells (GL7^-^), MBCs (CD38^+^CD98^-^), and PCs (CD38^low^CD98^+^). Quantifications indicate absolute numbers of (A) total YFP+ cells or (B) YFP+ GC B cells, MBCs and PCs. (**C**) Histograms and quantifications show the expression of CD19, CD39 and CD95 in lung MBCs and PCs of influenza-infected Aid-EYFP mice. (**D**) Flow cytometry plots displaying YFP+ GC B cells, MBCs and PCs in lungs of Aid-EYFP mice 365 days post infection with influenza virus. Quantifications indicate absolute numbers of YFP+ GC B cells, MBCs and PCs. (**E**) Bar chart shows the PC:MBC ratio in lungs 365 days post infection with influenza virus. (**F**) Flow cytometry plots of mes LN cells in non-infected (NI) or Salmonella-infected AID-EYFP mice and treated as in Figure 1A. (**G**) Flow cytometry plots display the gating strategy of YFP^+^ GC B cells (GL7^+^), memory cells (GL7^-^), MBCs (CD38^+^CD98^-^), and PCs (CD38^low^CD98^+^). Quantifications indicate absolute numbers of (F) total YFP+ cells or (G) YFP+ GC B cells, MBCs and PCs. (**H**) Histograms and quantifications show the expression of CD19, CD39 and CD95 in MBCs and PCs in small (upper panel) and large (lower panel) intestines of Salmonella-infected Aid-EYFP mice. (**I**) Enumeration of LPS-specific antibody-secreting cells in the small (upper panel) and large (lower panel) intestine 250 days after *Salmonella* infection. (**J**) Quantification of LPS-specific immunoglobulin levels in fecal extracts 250 days after *Salmonella* infection. (**K**) Flow cytometry plots display YFP^+^ PCs in the bone marrow of Aid-EYFP mice infected with *Klebsiella* intranasally or orally (day 35). Bar charts show the quantification in percentage and cell numbers. (**L**) Enumeration of LPS-specific antibody-secreting cells in the bone marrow 35 days after *Klebsiella* infection intranasally or orally. All panels display quantification from one representative experiment out of three (mean ± s.e.m.). Each dot represents one mouse. t-test (A, F, J, K) and two-way ANOVA (B, D, G, I and L), t-test to calculate whether the PC/MBC ratio was < or > 1 (E): *p<0.05 and ****p<0.0001.

**Figure S2.**
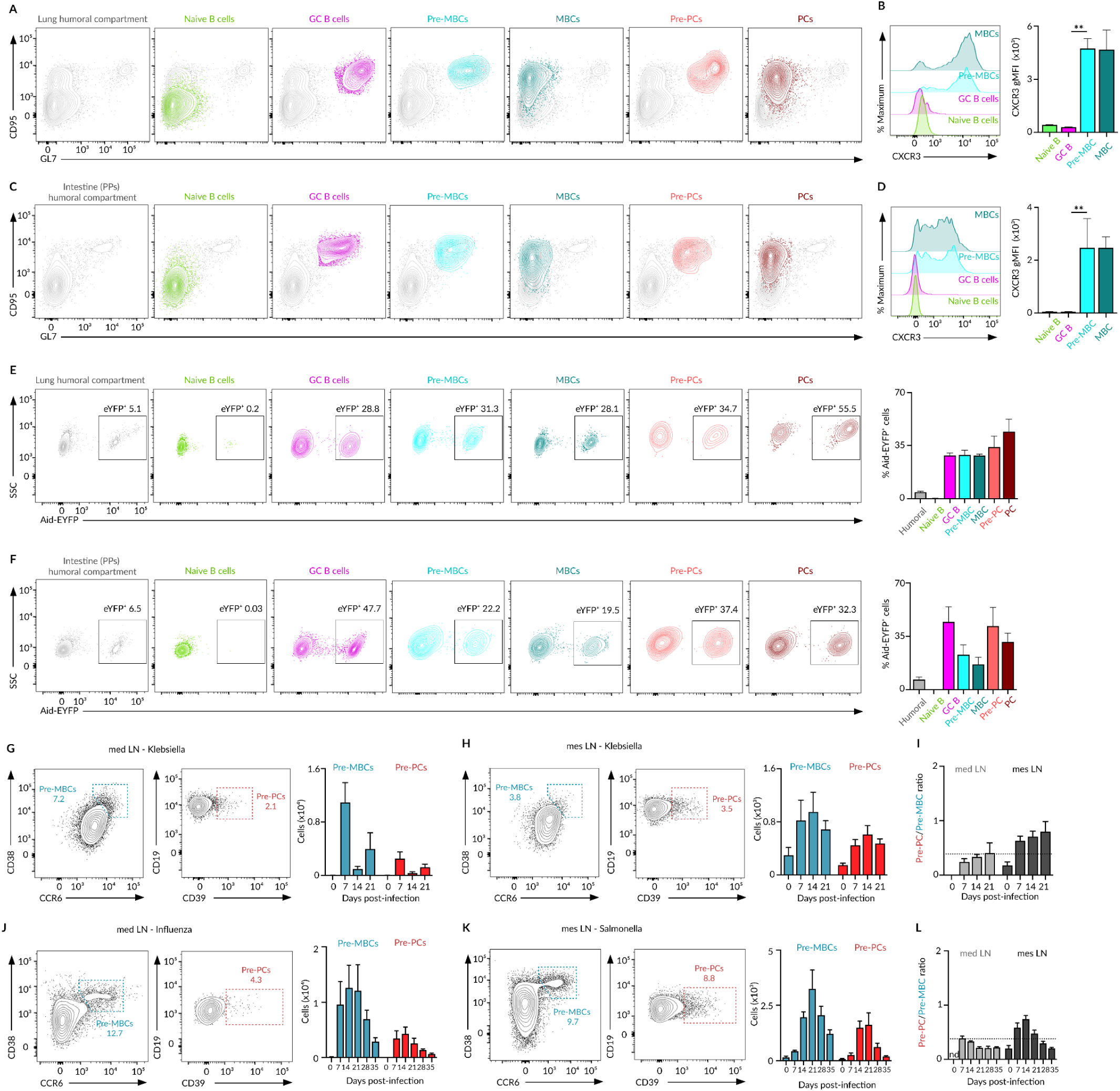
Characterization of mucosal Pre-MBC and Pre-PC compartments. (**A** and **C**) Contour plots showing the expression of CD95 and GL7 markers by naive B cells, GC B cells, pre-MBCs, MBCs, pre-PCs and PCs populations in the (A) lungs or (C) Peyer’s patches of Influenza or Salmonella-infected Aid-EYFP mice. (**B** and **D**) Histograms and quantification of the expression of CXCR3 among Naive B cells, GC B cells, pre-MBCs and MBCs populations in the (B) lungs or (D) Peyer’s patches of influenza or Salmonella infected mice. (**E** and **F**) Flow cytometry plots and quantification of eYFP+ cells within Naive B cells, GC B cells, pre-MBCs, MBCs, pre-PCs and PCs populations in the (**E**) lungs or (**F**) Peyer’s patches of influenza or Salmonella infected mice. (**G, H, J** and **K**) Flow cytometry plots and quantification of pre-MBCs (CD38^+^CCR6^+^) and pre-PCs (CD19^+^CD39^+^) within the GC reaction (CD95^+^GL7^+^) in (G and J) med lymph nodes and (H and K) mes lymph nodes after (G and H) Klebsiella or (J and K) Influenza or Salmonella infection. (**I** and **L**) Bar charts showing the prePC:preMBC ratio in med and mes lymph nodes after (I) Klebsiella or (L) Influenza or Salmonella infection.

**Figure S3.**
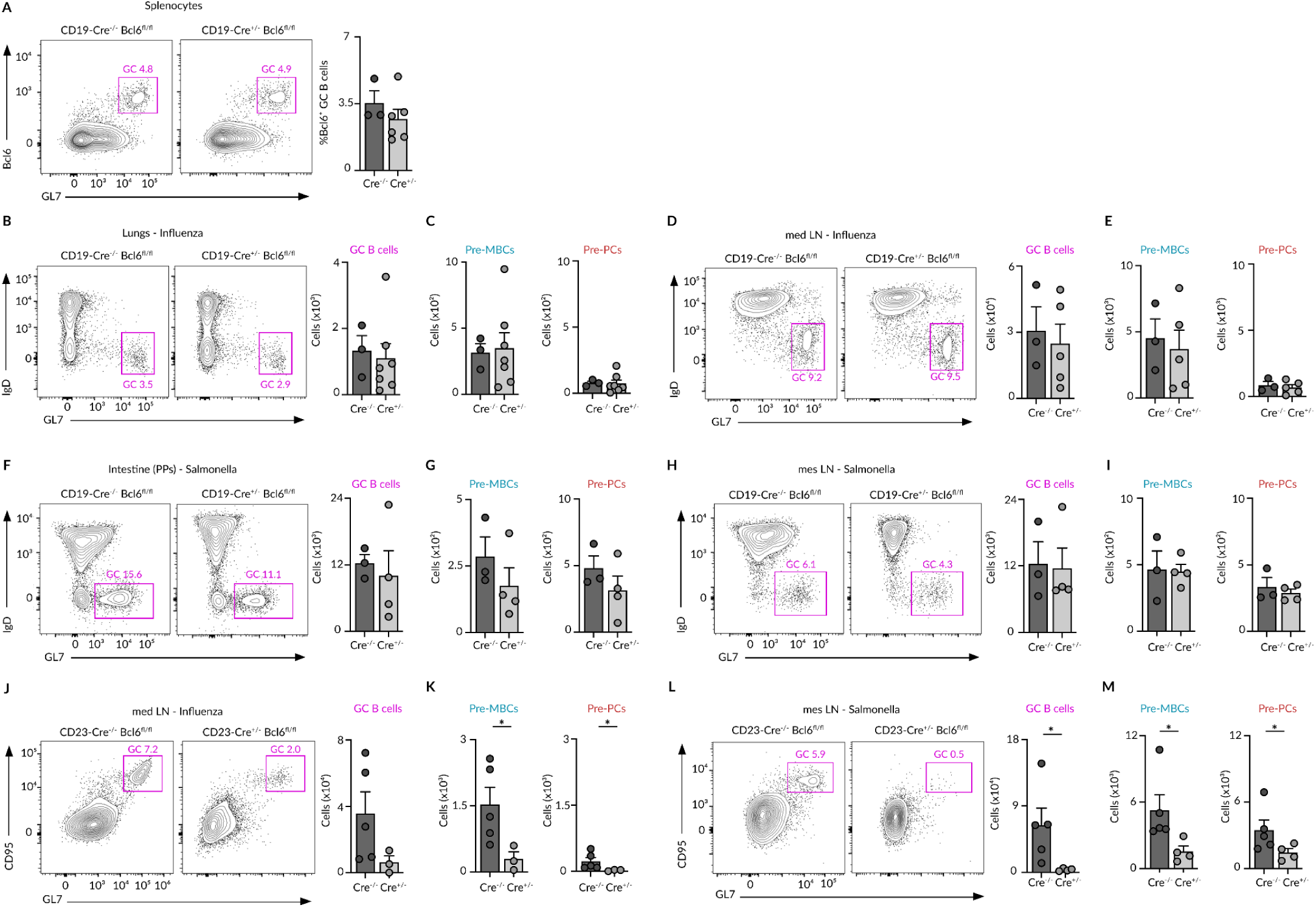
GC origin of Pre-MBC and Pre-PCs. (**A**) Contour plots and quantification of Bcl6-expressing GC B cells in the spleen of CD19Cre-Bcl6^fl/fl^ mice after influenza infection. (**B** and **D**) Contour plots and quantification of GC B cells (IgD^-^GL7^+^) in (B) lungs and (D) med lymph nodes of CD19Cre-Bcl6^fl/fl^ mice after influenza infection. (**C** and **E**) Quantification of pre-MBCs and pre-PCs in (C) lungs and (E) med lymph nodes of CD19Cre-Bcl6^fl/fl^ mice after influenza infection. (**F** and **H**) Contour plots and quantification of GC B cells (IgD^-^GL7^+^) in (F) Peyer’s patches and (H) mes lymph nodes of CD19Cre-Bcl6^fl/fl^ mice after Salmonella infection. (**G** and **I**) Quantification of pre-MBCs and pre-PCs in (G) Peyer’s patches and (I) mes lymph nodes of CD19-cre Bcl6^fl/fl^ mice after Salmonella infection. (**J** and **L**) Contour plots and quantification of GC B cells (CD95^+^GL7^+^) in (J) med and (L) mes lymph nodes of CD23Cre-Bcl6^fl/fl^ mice after influenza or Salmonella infection. (**K** and **M**) Quantification of pre-MBCs and pre-PCs in (K) med and (M) mes lymph nodes of CD23Cre-Bcl6^fl/fl^ mice after influenza or Salmonella infection. All panels from CD23Cre-Bcl6^fl/fl^ mice display quantification from one representative experiment out of three (mean ± s.e.m.). Each dot represents one mouse. t-test (**A**-**M**): *p<0.05.

**Figure S4.**
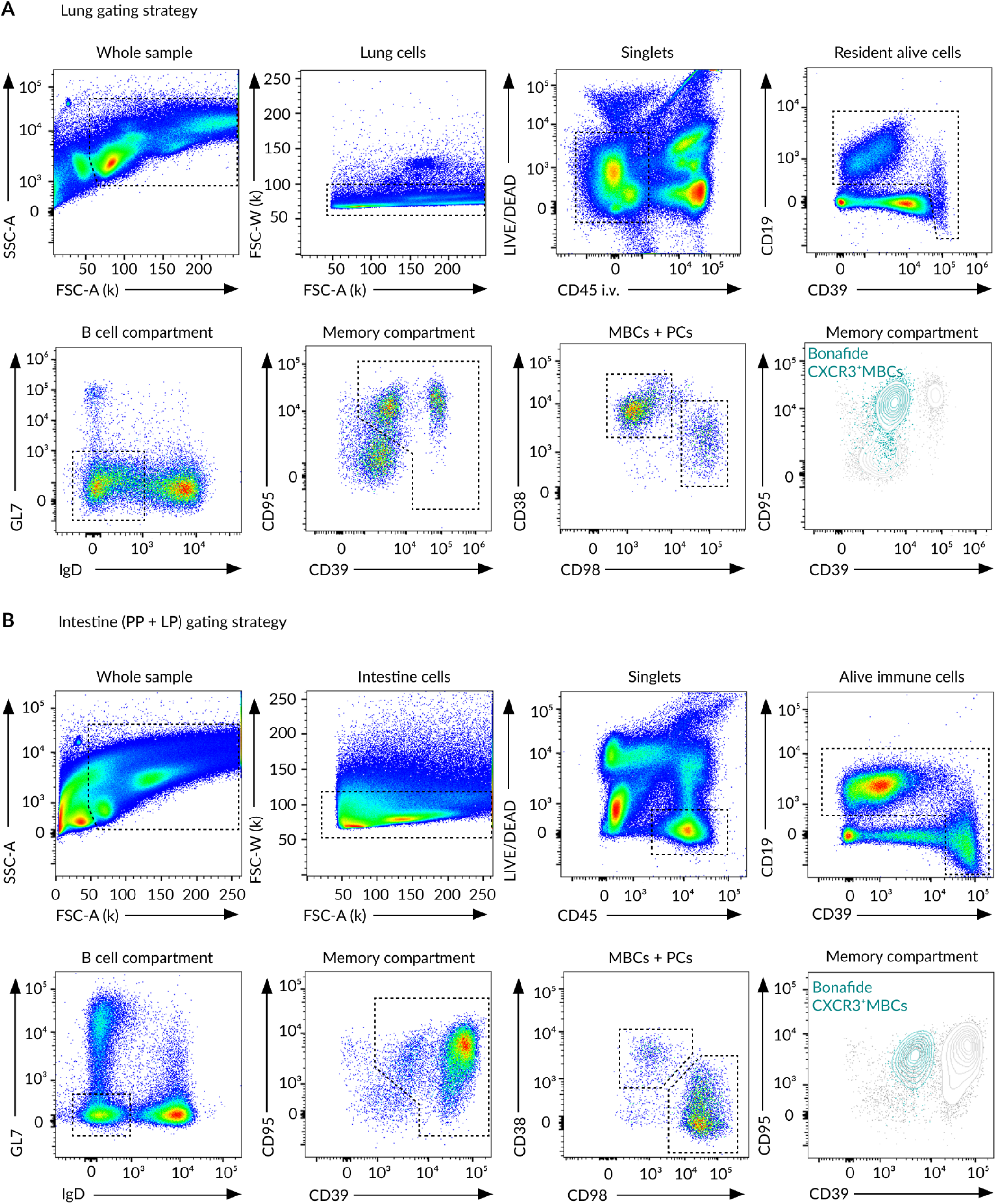
Gating strategy used to identify MBCs and PCs in non Aid-EYFP mice. (**A** and **B**) Flow cytometry plots showing the gating strategy for MBCs and PCs in (**A**) lungs and (**B**) intestine after infection.

**Figure S5.**
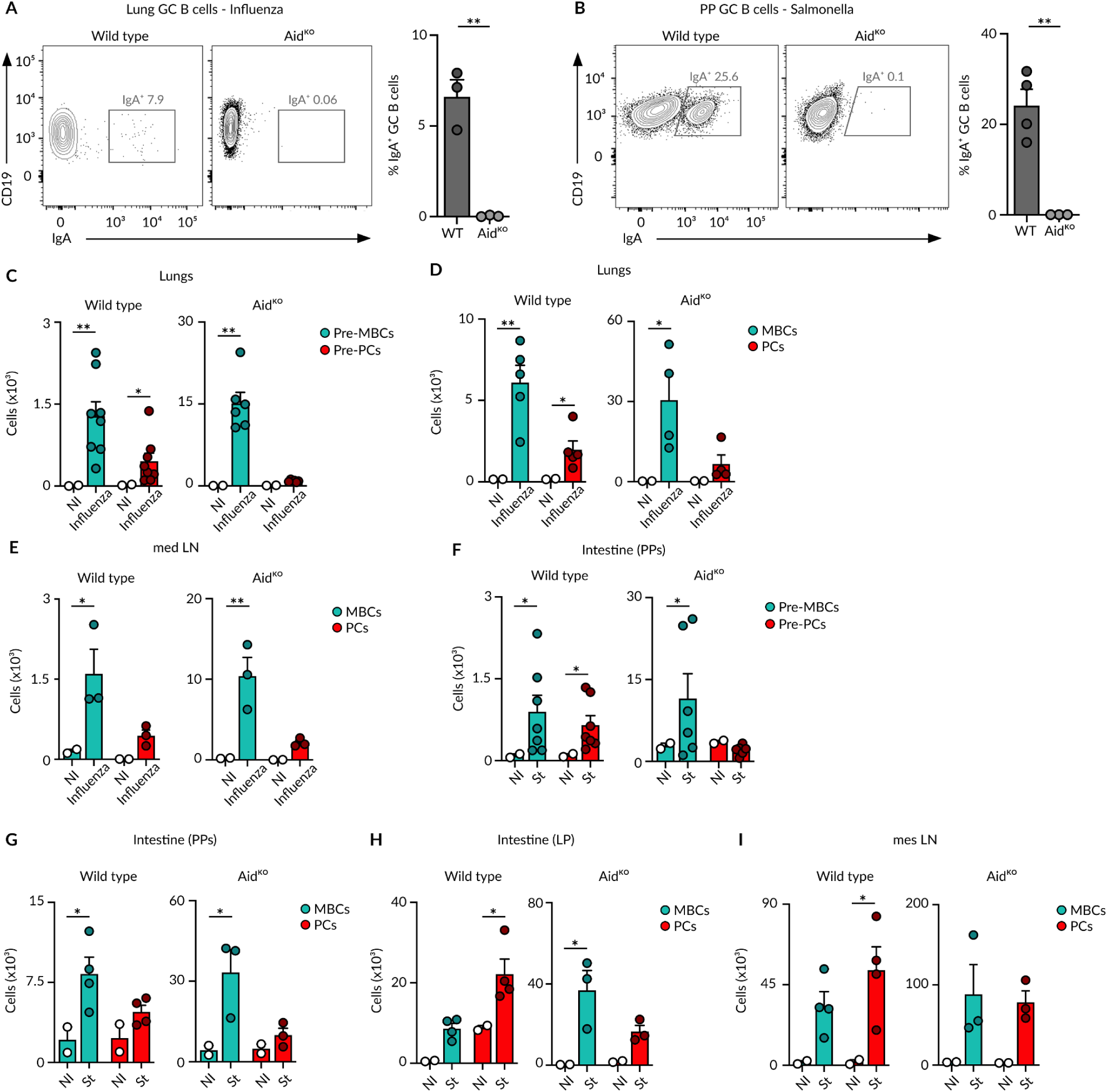
Characterization of mucosal pre-memory and memory compartments in Aid-KO mice. (**A** and **B**) Flow cytometry plots display IgA expression by GC B cells in (A) lung of influenza-infected and (B) Peyer’s patches of *Salmonella*-infected wild type and *Aid* KO mice. Bar charts show IgA^+^ B cells in the GC population. (**C** and **F**) Quantification of pre-MBC and pre-PC numbers in (C) lungs of influenza-infected and (F) Peyer’s patches of *Salmonella*-infected wild type and *Aid*-KO mice. (**D** and **E**) Quantification of MBC and PC numbers in (D) lungs and (E) med lymph nodes of influenza-infected wild type and *Aid*-KO mice. (**G, H** and **I**) Quantification of MBC and PC numbers in (G) Peyer’s patches, (H) lamina propria and (F) mes lymph nodes of Salmonella-infected wild type and *Aid*-KO mice. All panels display quantification from one representative experiment out of three (mean ± s.e.m.). Each dot represents one mouse. t-test (A and B) and two-way ANOVA (C-I): *p<0.05 and **p<0.01.

**Figure S6.**
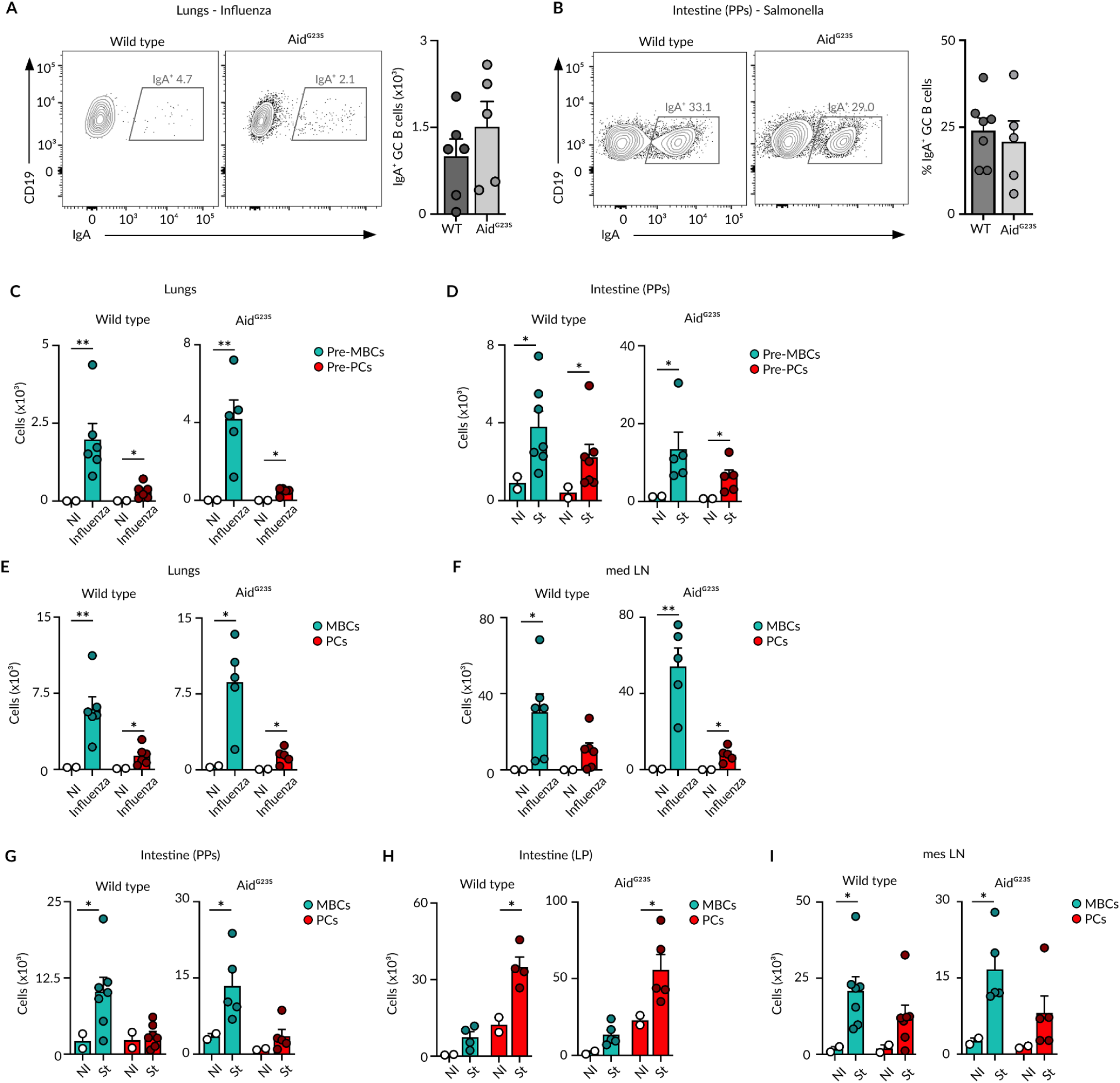
Characterization of mucosal pre-memory and memory compartments in Aid(G23S) mice. (**A** and **B**) Flow cytometry plots display IgA expression by GC B cells in (A) lung of influenza-infected and (B) Peyer’s patches of *Salmonella*-infected wild type and *Aid*(G23S) mice. Bar charts show IgA^+^ B cells in the GC population. (**C** and **D**) Quantification of pre-MBC and pre-PC numbers in (C) lungs of influenza-infected and (D) Peyer’s patches of *Salmonella*-infected wild type and *Aid*(G23S) mice. (**E** and **F**) Quantification of MBC and PC numbers in (E) lungs and (F) med lymph nodes of influenza-infected wild type and *Aid*(G23S) mice. (**G, H** and **I**) Quantification of MBC and PC numbers in (G) Peyer’s patches, (H) lamina propria and (F) mes lymph nodes of Salmonella-infected wild type and *Aid*(G23S) mice. All panels display quantification from one representative experiment out of three (mean ± s.e.m.). Each dot represents one mouse. t-test (A and B) and two-way ANOVA (C-I): *p<0.05 and **p<0.01.

**Figure S7.**
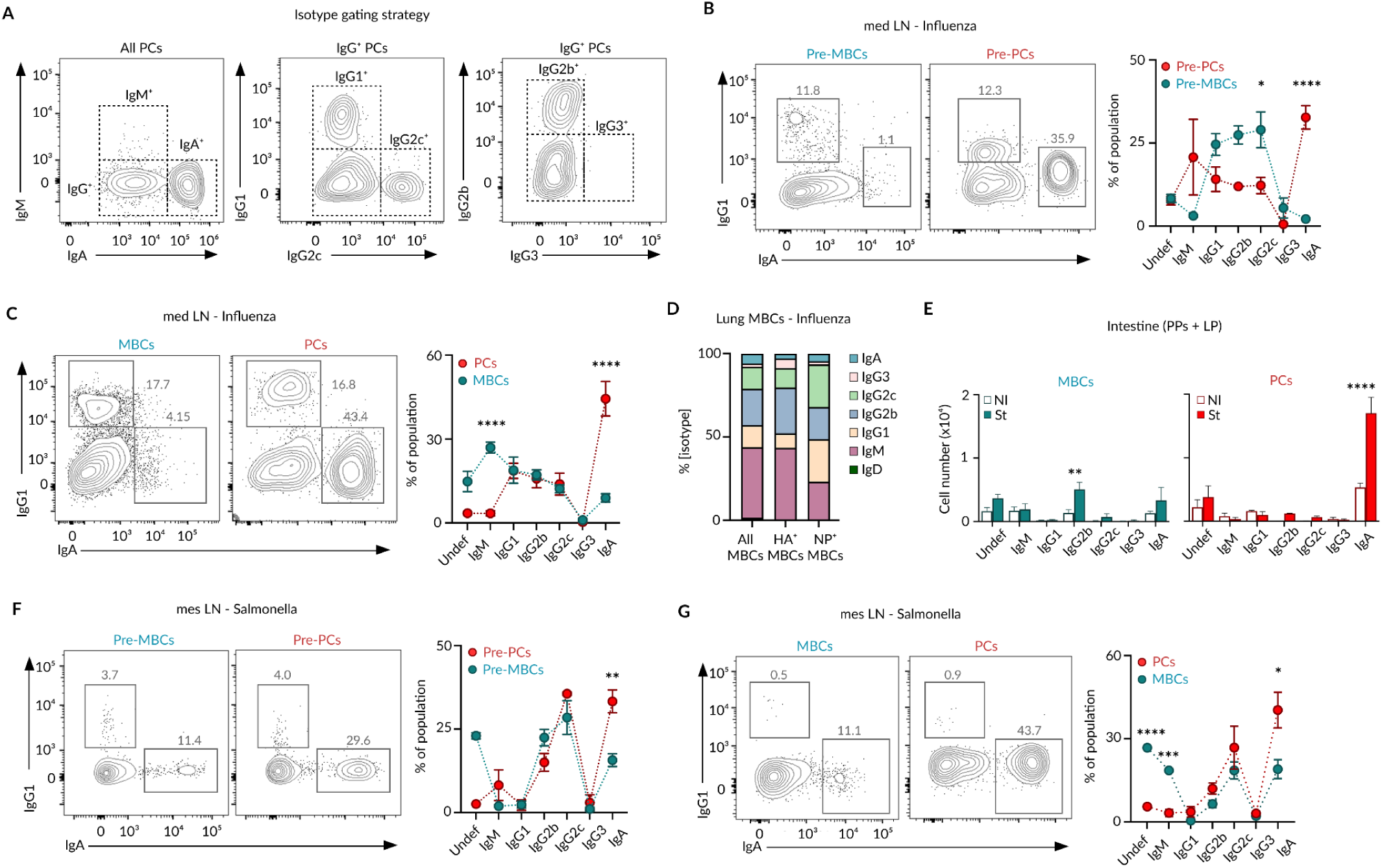
Characterization of isotype distribution by mucosal pre-memory and memory compartments. (**A**) Contour plots showing the gating strategy used to identify isotype^+^ B cells. (**B** and **C**) Contour plots and quantification of isotype distribution in (B) pre-MBCs and pre-PCs and (C) MBCs and PCs in med lymph nodes of influenza-infected mice. (**D**) BCR isotype distribution in total, HA^+^ or NP^+^ MBCs in the lungs of influenza-infected mice (data from Gregoire et al. 2022). (**E**) Isotype distribution in intestinal MBCs and PCs in non-infected or Salmonella-infected mice. (**F** and **G**) Contour plots and quantification of isotype distribution in (F) pre-MBCs and pre-PCs and (G) MBCs and PCs in mes lymph nodes of Salmonella-infected mice. All panels display quantification from one representative experiment out of three (mean ± s.e.m.). Two-way ANOVA (B, C, E, F and G): *p<0.05, **p<0.01, ***p<0.001 and ****p<0.0001.

**Figure S8.**
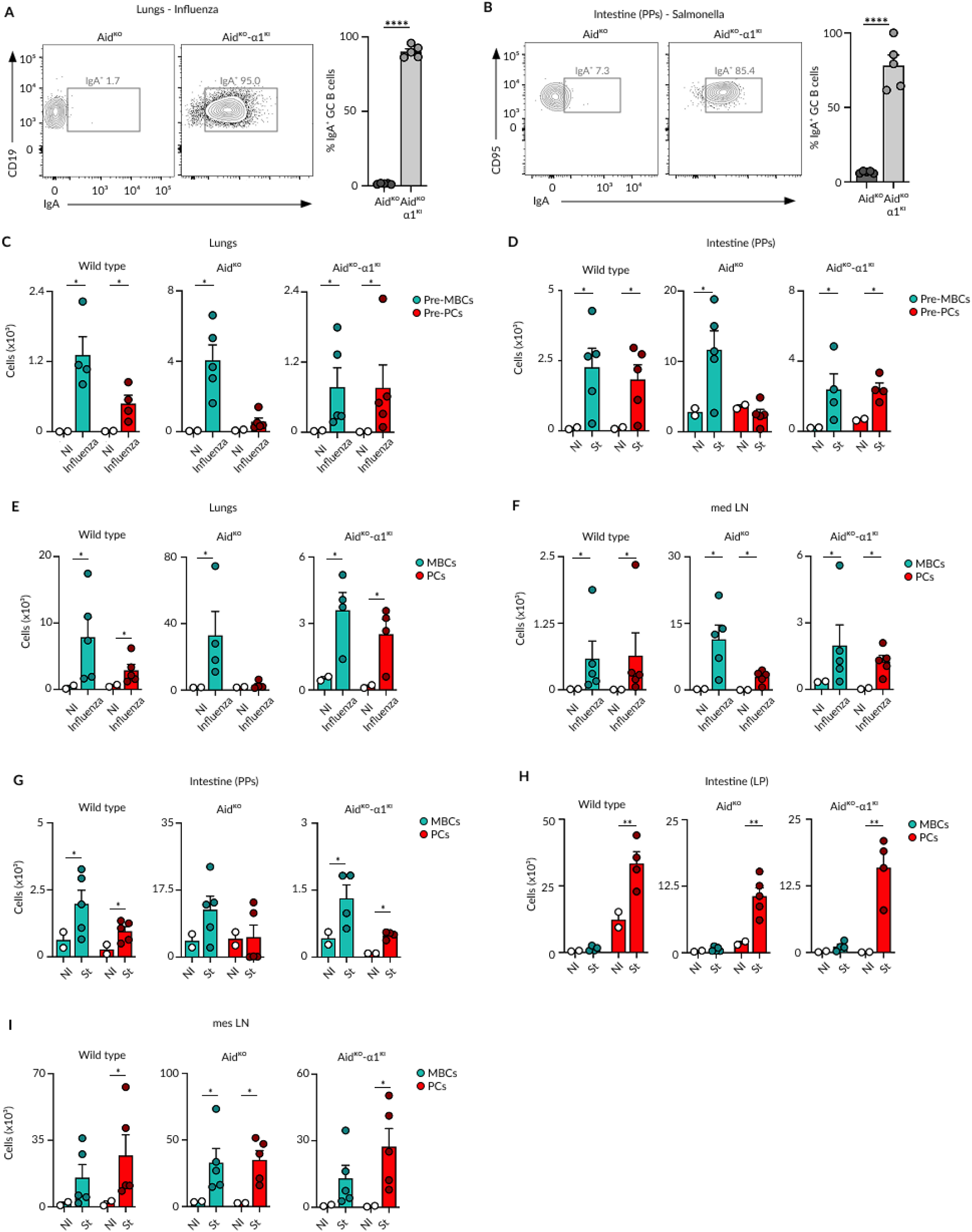
Characterization of mucosal pre-memory and memory compartments in AidKO-α1KI mice. (**A** and **B**) Flow cytometry plots display IgA expression by GC B cells in (A) lung of influenza-infected and (B) Peyer’s patches of *Salmonella*-infected *Aid*-KO and *Aid*-KO α1KI mice. Bar charts show IgA^+^ B cells in the GC population. (**C** and **D**) Quantification of pre-MBC and pre-PC numbers in (C) lungs of influenza-infected and (D) Peyer’s patches of *Salmonella*-infected wild type, *Aid*-KO and *Aid*-KO α1KI mice. (**E** and **F**) Quantification of MBC and PC numbers in (E) lungs and (F) med lymph nodes of influenza-infected wild type, *Aid*-KO and *Aid*-KO α1KI mice. (**G, H** and **I**) Quantification of MBC and PC numbers in (G) Peyer’s patches, (H) lamina propria and (F) mes lymph nodes of Salmonella-infected wild type, *Aid*-KO and *Aid*-KO α1KI mice. All panels display quantification from one representative experiment out of three (mean ± s.e.m.). Each dot represents one mouse. t-test (A and B) and two-way ANOVA (C-I): *p<0.05, **p<0.01 and ****p<0.0001.

**Figure S9.**
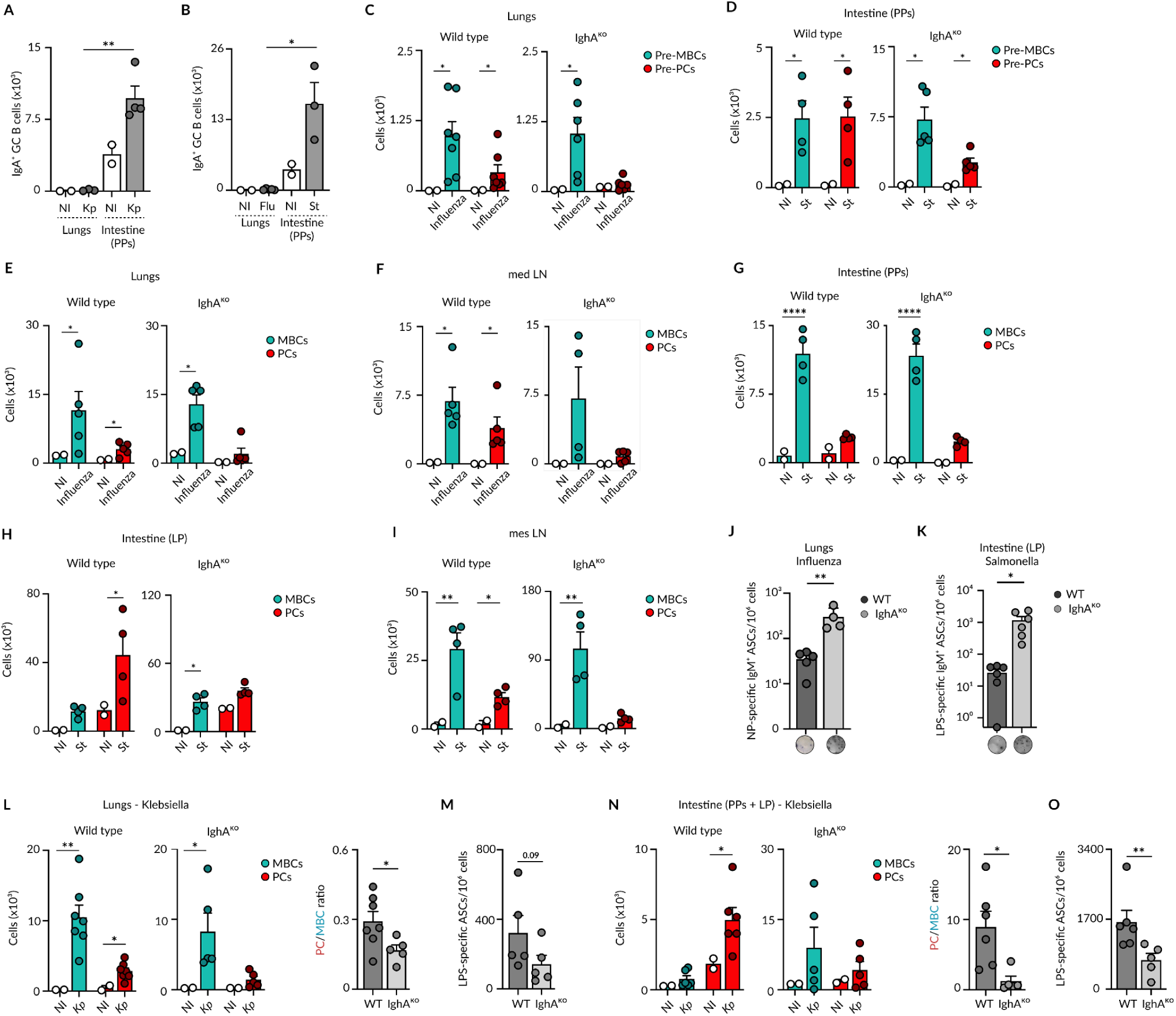
Characterization of mucosal pre-memory and memory compartments in IghA-KO mice. (**A** and **B**) Bar charts show the quantification of IgA^+^ B cells in the GC population in lungs or intestine after (A) *Klebsiella* or (B) influenza or *Salmonella* infection. (**C** and **D**) Quantification of pre-MBC and pre-PC numbers in (C) lungs of influenza-infected and (D) Peyer’s patches of *Salmonella*-infected wild type and *IghA-*KO mice. (**E** and **F**) Quantification of MBC and PC numbers in (E) lungs and (F) med lymph nodes of influenza-infected wild type and *IghA-*KO mice. (**G, H** and **I**) Quantification of MBC and PC numbers in (G) Peyer’s patches, (H) lamina propria and (F) mes lymph nodes of Salmonella-infected wild type and *IghA-*KO mice. (**J**) Enumeration of lung NP-specific IgM^+^ antibody-secreting cells in influenza-infected wild type and *Igha* KO mice by ELISpot. (**K**) Enumeration of intestinal LPS-specific IgM^+^ antibody-secreting cells in *Salmonella*-infected wild type and *Igha* KO mice by ELISpot. (**L** and **N**) Quantification of MBC and PC numbers in (L) lungs and (N) intestine of *Klebsiella*-infected wild type and *Igha* KO mice. The right bar charts show PC:MBC ratio in (L) lungs and (N) intestine of wild type and *Igha*-KO mice infected with *Klebsiella.* (**M** and **O**) Enumeration of (M) lung or (O) intestinal LPS-specific antibody secreting cells by ELISpot. All panels display quantification from one representative experiment out of three (mean ± s.e.m.). Each dot represents one mouse. t-test (**J**, **K**, **L**, **M, N** and **O**) and two-way ANOVA (**A**-**I, L** and **N**): *p<0.05, **p<0.01 and ****p<0.0001.

**Figure S10.**
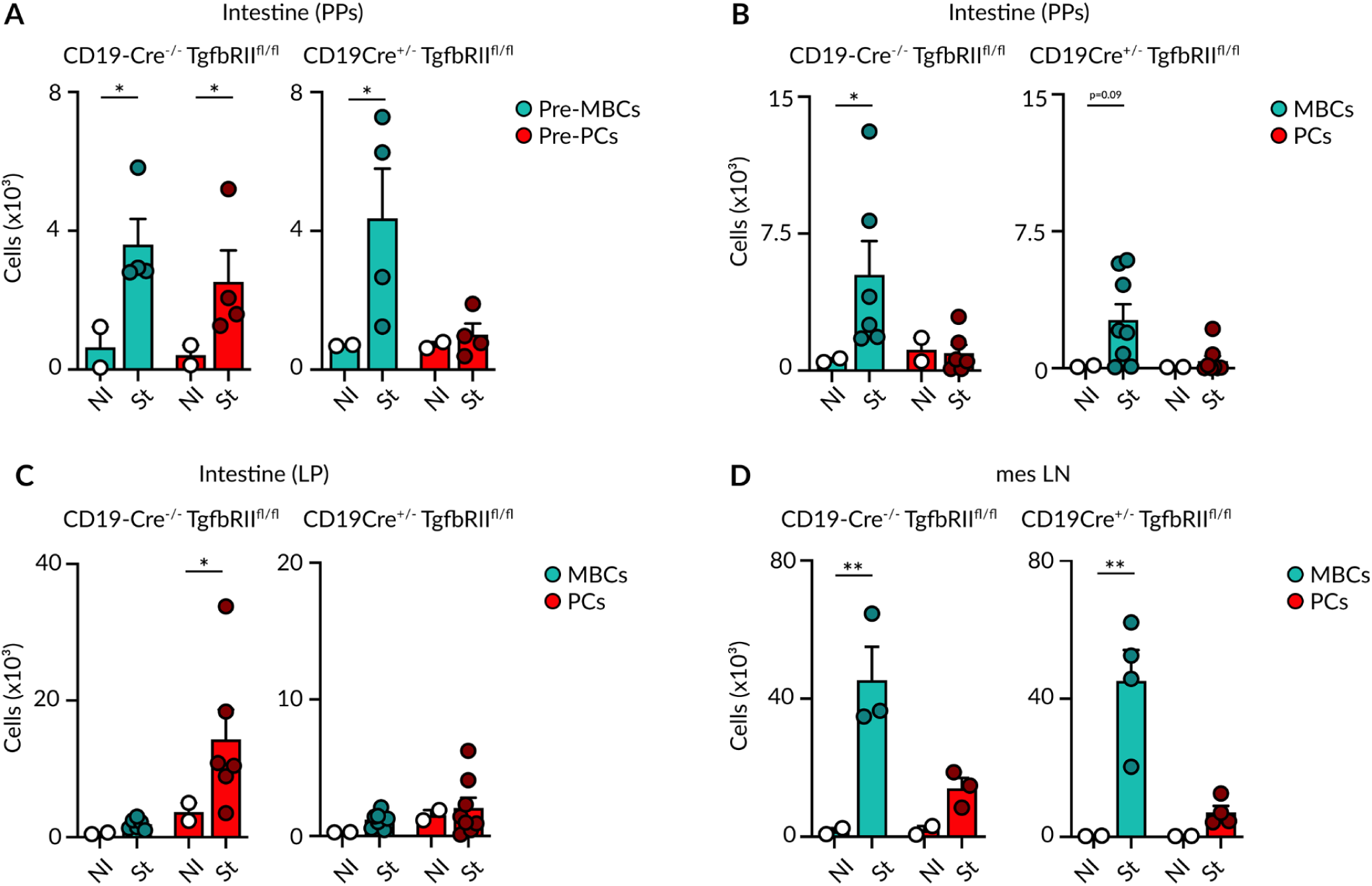
Characterization of mucosal pre-memory and memory compartments in CD19Cre-TgfbRII^fl/fl^ mice. **(A)** Quantification of pre-MBC and pre-PC numbers in Peyer’s patches of *Salmonella*-infected CD19Cre-TgfbRII^fl/fl^ mice. (**B, C** and **D**) Quantification of MBC and PC numbers in (B) Peyer’s patches, (C) lamina propria and (D) mes lymph nodes of Salmonella-infected CD19Cre-TgfbRII^fl/fl^ mice. All panels display quantification from one representative experiment out of three (mean ± s.e.m.). Each dot represents one mouse. Two-way ANOVA (A-D): *p<0.05 and **p<0.01.

**Figure S11.**
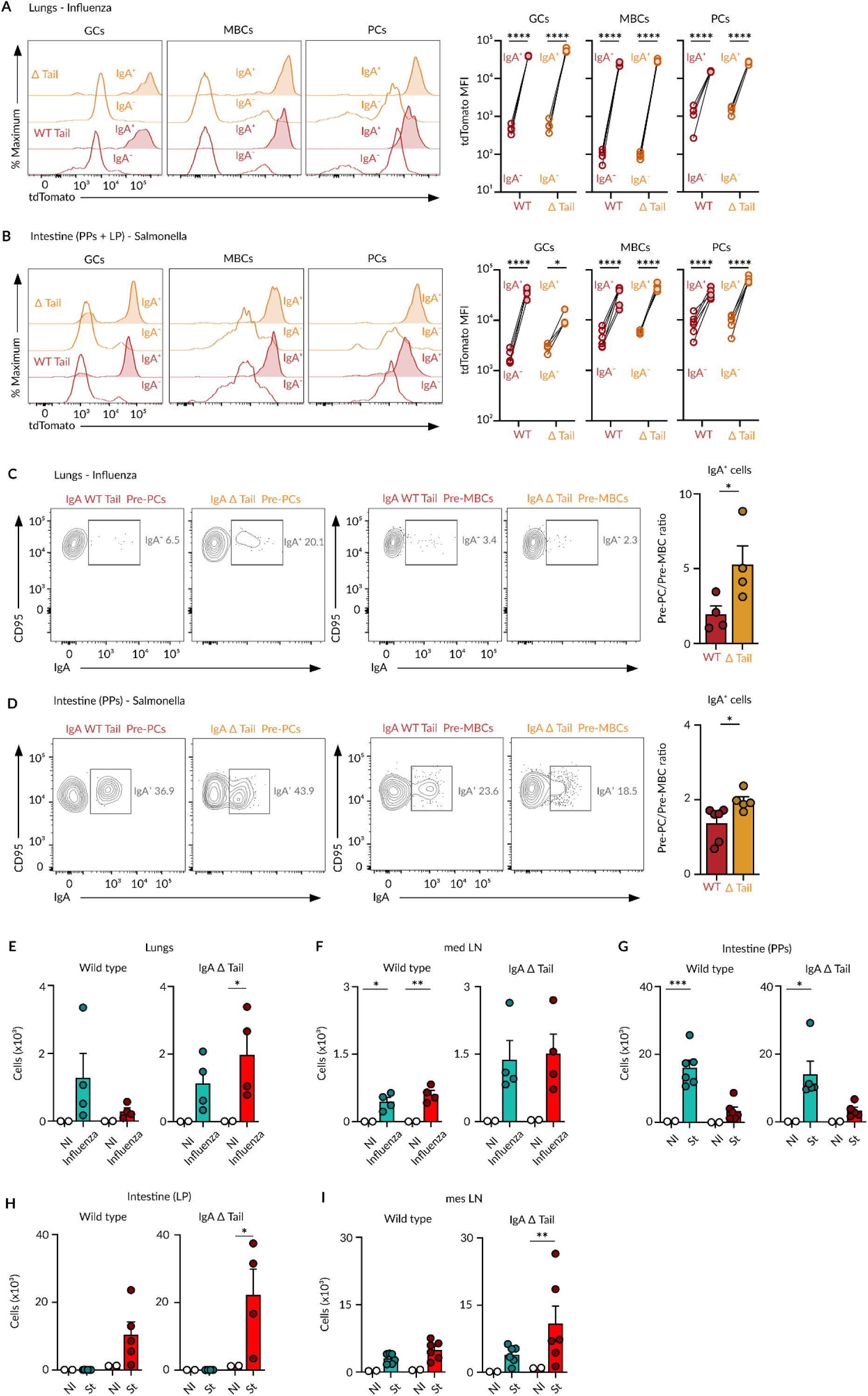
Characterization of mucosal pre-memory and memory compartments in Igha-tdTom and Igha-Δtail-tdTom mice. (**A** and **B**) Histograms displaying tdTomato expression by IgA^-^ and IgA^+^ GCs, MBCs and PCs in (A) lungs and (B) intestine of c infected with (A) influenza virus or (B) *Salmonella*, respectively (day 35). Quantifications show tdTom MFI for all populations. (**C** and **D**) Contour plots displaying IgA^+^ cells in pre-PCs and pre-MBCs populations in (**C**) lungs and (**D**) intestines of Igha-tdTom and Igha-Δtail-tdTom mice infected with influenza virus or *Salmonella*, respectively. Quantifications show IgA^+^ pre-PCs: IgA^+^ pre-MBCs ratios. (**E** and **F**) Quantification of MBC and PC numbers in the (E) lungs and (F) med LN of Influenza-infected Igha-tdTom and Igha-Δtail-tdTom mice. (**G**-**I**) Quantification of MBC and PC numbers in the (G) PPs, (H) Lamina Propria and (I) mes LN of *Salmonella* infected Igha-tdTom and Igha-Δtail-tdTom mice. In all panels, quantifications display one representative experiment out of three (mean ± s.e.m.). Each dot represents one mouse. Two-way ANOVA: *p<0.05, **p<0.01, ***p<0.001 and ****p<0.0001.

**Figure S12.**
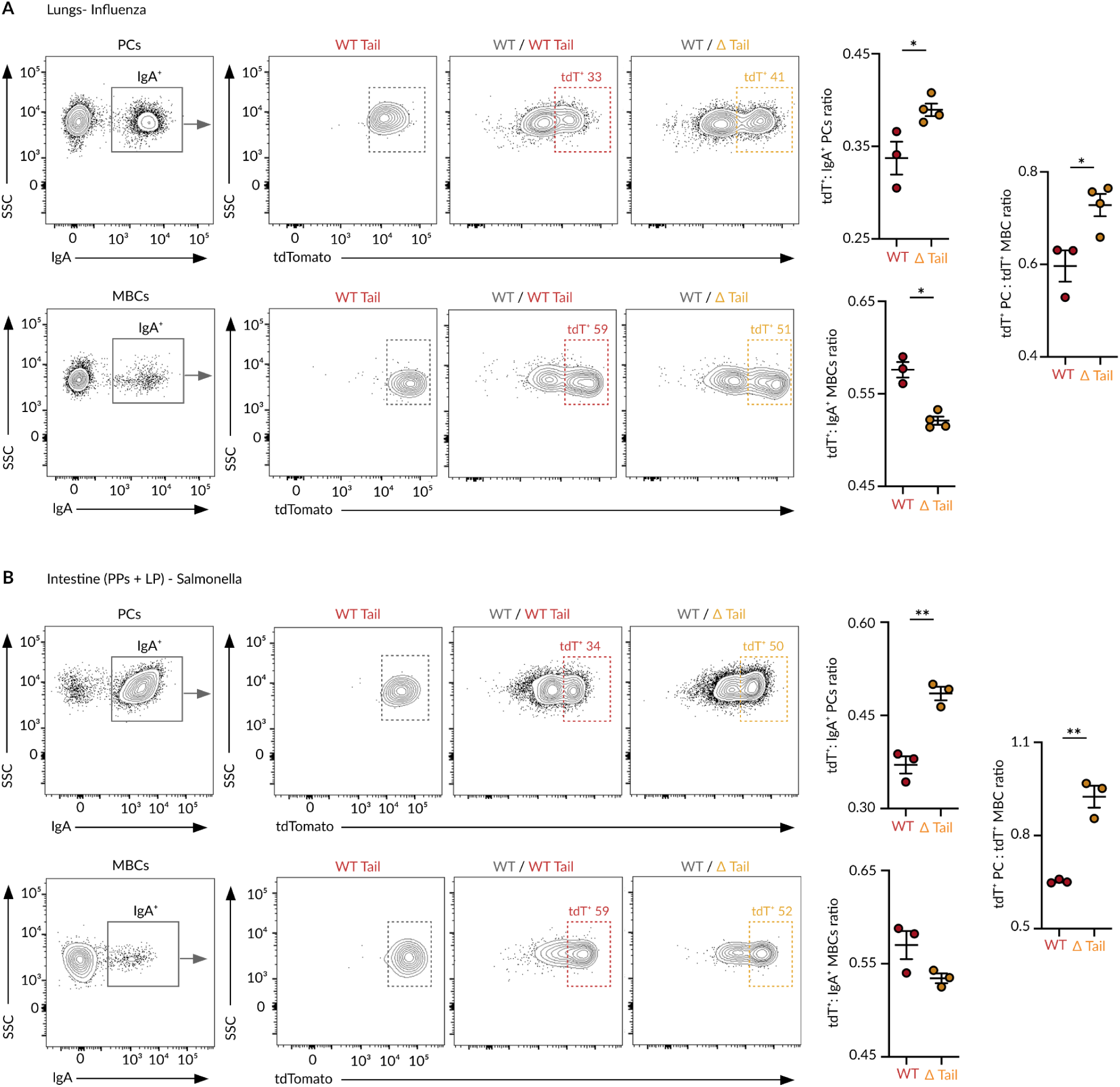
Mucosal B cell memory instruction by the IgA cytosolic tail in a competitive system. (**A-B**) Contour plots displaying tdTomato expression by IgA^+^ MBCs and PCs in (A) lungs and (B) intestine of Igha-tdTom+/+, Igha-tdTom+/- and Igha-Δtail-tdTom+/- mice infected with (A) influenza virus or (B) Salmonella, respectively (day 35). Quantifications show tdT+: IgA+ ratios for PCs and MBCs. In all panels, quantifications display one representative experiment, mean ± s.e.m. Each dot represents one mouse. t test: *p<0.05 and **p<0.01.

## Methods

### Mice

8-week old wild-type SPF C57BL/6 and BALB/cJRj mice were obtained from Janvier Labs. Aicda-CreERT2 mice were obtained from Claude-Agnès Reynaud and Jean Claude Weill, Institut Necker Enfants Malades, France. Rosa26-EYFP mice were obtained from Jackson Laboratories, USA. CD19-Cre mice were obtained from Sandrine Roulland, Centre d’Immunologie de Marseille Luminy, France. Igha-KO mice were obtained from Emma Slack, Institute of Food, Nutrition and Health, Switzerland. Tgfbr2^fl/fl^ mice were obtained from Julien Marie, Cancer Research Center of Lyon, France. Aid(G23S) mice were obtained from Tasuku Honjo, Center for Cancer Immunotherapy and Immunobiology, Japan. α1KI mice were obtained from Michel Cogné, Laboratoire d’Immunologie, Université de Limoges, France. CD23cre Bcl6^fl/fl^ mice were obtained from Michelle Linterman, Babraham Institute, UK. Aicda-Cre^ERT2^ mice were further crossed with Rosa26-EYFP mice. IgA-tdT mice (C57BL/6NRj-Igha^tm1Ciphe^) and Igha-Δtail-tdTom (C57BL/6NRj-Igha^em2Ciphe^) were generated in CIPHE (Centre d’Immunophénomique). Igha-tdTom mice were generated by homologous recombination in embryonic stem cells. Igha-Δtail-tdTom were obtained by deletion of Igha tail by electroporation of CRISPR/Cas9 complex and single-stranded donor DNA into homozygous zygotes for the Igha-tdT mice. Germ-free C57BL/6 mice were purchased from TAAM-CNRS (Orléans, France). Both germ-free and SPF mice were transferred to the same conventional barrier facility for infection, allowing us to compare the impact of their initial gut environment on subsequent infection while maintaining identical experimental conditions.

Mice were bred and maintained at the animal facilities of the Centre d’Immunologie de Marseille Luminy and Ciphe. Up to five mice per cage were housed under a standard 12 hr light/dark cycle, with room temperature at 22°C (19°C-23°C change). They were fed with autoclaved standard pellet chow or tamoxifen pellet chow when indicated, and reverse osmosis water. All cages contained 5 mm of aspen chip and tissue wipes for bedding and a mouse house for environmental enrichment. Mice were used at the age of 8 to 12 weeks and littermates (males or females) were randomly assigned to experimental groups. Generally between 4 to 5 mice were used per experimental group. Experimental procedures were conducted in accordance with French and European guidelines for animal care under the permission number 16708-2018091116493528 and 43129-202304171838427 following review and approval by the local animal ethics committee in Marseille.

### Infections and injections

For intranasally infections, mice were anesthetized i.p. with Ketamine/Xylazine (100 mg/kg body) and intranasally infected with 5 PFU of Influenza virus A/Puerto Rico/8/1934 (PR8) H1N1 strain, 5x10^4^ PFU of Influenza virus A/X-31 H3N2, 10^7^ CFU of *Klebsiella pneumoniae* 43816 strain or 5x10^3^ CFU of *Klebsiella pneumoniae* 52145 strain in 20 μl of PBS. Influenza virus was amplified on MDCK cells. Purification of viral particles was performed in a sucrose 30% cushion at 25,000 RPM for 2 hours in an SW32Ti rotor. For Aid-EYFP mice, tamoxifen (Cayman chemical) was resuspended in corn oil (Sigma), sonicated and given by oral gavage at 5mg/100μl. For gastrointestinal infections, mice were orally gavaged with 10^10^ CFU of *Salmonella typhimurium* aroA SL3261, 5x10^6^ CFU *Salmonella* WT or 10^10^ CFU of *Klebsiella pneumonia*e 43816 strain in 100 ul of PBS with a reusable feeding needle. *Salmonella typhimurium aro*A SL3261 and WT strains were obtained from David Holden, Imperial College London, UK. *Klebsiella pneumoniae* was obtained from Régis Tournebize, Institut Pasteur, France. For *in vivo* labeling of immune cells in circulation, 3 μg of anti-CD45 antibody was administered i.v 5 minutes before sacrifice.

### Flow Cytometry

For isolation of lamina propria cells, the intestine was divided into small and large sections. Peyer’s patches and caecum were excised from small and large intestine respectively, and processed afterwards. Small and large intestines were opened longitudinally and incubated in HBSS 2mM EDTA twice (1x 20 min and 1 x 30 min) at 37°C to remove epithelial cells. Then, cells were washed twice with HBSS Ca^+2^Mg^+2^ and incubated for 25 (small intestine) or 35 min (large intestine) in RPMI 1640 with 10% FCS, 1mg/ml of collagenase IV (Sigma), DNAse I 30ug/ml. The reaction was stopped with 1:100 of 0.5M EDTA, filtered consecutively through 100- and 70-µm mesh and the cell suspension was finally purified by density gradient centrifugation with 40–80% Percoll (GE Healthcare). Single cell suspensions of Peyer’s patches, mediastinal or mesenteric lymph nodes and spleens were obtained by pressing organs through a 70μm nylon mesh cell strainer with a plastic plunge in PBS 2%FCS 5mM EDTA. For lungs, single cell suspensions were obtained with the mouse lung dissociation kit (Miltenyi) according to manufacturer instructions and lymphocytes were enriched by using 40-80% Percoll gradient. For spleen and lungs, cell suspensions were further incubated with a red blood cell lysis buffer for 5 minutes. To block nonspecific antibody binding, cell suspensions were incubated with hybridoma supernatant 2.4G2 diluted 1/4 in PBS 2%FCS 5mM EDTA and when indicated with LIVE/DEAD Fixable Blue Dead Cell Stain Kit (Thermo Fisher Scientific) to exclude dead cells, for 20 minutes on ice. For cell death assay, 1 x 10⁶ cells were stained with NucView® Caspase-3 Enzyme Substrates (Biotium, 10402) during 30 minutes at room temperature. Regarding ROS measurement, 1 x 10^7^ cells were cultured overnight with 400µL of complete RPMI medium, containing or not anti-IgA (SouthernBiotech, 10µg/mL). Cells were then stained with 100µL of MitoSox green (ThermoFischer, 5µM) diluted in FACS buffer, during 30 minutes at 37°C. For labeling of surface markers, cells were stained for 20 minutes on ice with the indicated anti-mouse antibodies. For labelling of BCR isotypes, cell suspensions were fixed 30 minutes with PFA 4% and intracellular staining was performed with Cytofix/Cytoperm Fixation/Permeabilization Kit (BD Biosciences) following manufacturer protocol. The antibodies used to identify BCR isotypes are listed in the table below. Finally, cells were either resuspended in 400μl of PBS 2%FCS 2mM EDTA together with the addition of beads (CountBright™ Absolute Counting Beads, Invitrogen) and analyzed on Fortessa-X20/Symphony/LSRIIUV cytometers (BD Biosciences) or in complete RPMI medium (FCS 10%, Penicillin 100µg/ml, Streptomycin 10μL/ml, Sodium Pyruvate 1mM, 50μM 2-mercaptoethanol) and used for cell sorting on a FACSAria II (BD, Bioscience). For non-fixed cells, a mix of counting beads and DAPI (0.1 μg/ml final concentration) was added right before passing samples on cytometers. Data was analyzed using FlowJo (TreeStar).

| Antibody | Source | Catalog |
| --- | --- | --- |
| Alexa Fluor 488 anti-mouse IgG2b Antibody, Clone RMG2b-1 | Biolegend | 406718 |
| Alexa Fluor 647 Goat Anti-Mouse IgA, polyclonal | Southern BT | 1040-31 |
| Alexa Fluor 647 Rat Anti-Mouse Bcl-6, clone K112-91 | BD Horizon | 561525 |
| Alexa Fluor 647 Goat anti-Mouse IgG2a Cross-Adsorbed Secondary Antibody, polyclonal | ThermoFisher | A-21241 |
| Alexa Fluor 647 Rat Anti-Mouse/Human GL7 Antigen, Clone GL7 | Biolegend | 144606 |
| APC CD90.1 (Thy-1.1), Clone HIS51 | ThermoFisher | 17-0900-82 |
| APC/Cy7 Rat Anti-Mouse CD19, Clone 6D5 | Biolegend | 115530 |
| BUV395 Rat Anti-Mouse CD45R/B220, clone RA3-6B2 | BD Horizon | 563793 |
| BUV395 Rat Anti-Mouse CD38, Clone 90/CD38 | BD Horizon | 740245 |
| BUV737 Rat Anti-Mouse CD98, Clone H202-141 | BD | 749412 |
| BUV737 Rat anti-Mouse/Human CD45R (B220), Clone RA3-6B2 | BD Horizon | 612839 |
| BV421 Armenian Hamster Anti-Mouse CD183 (CXCR3), Clone CXCR3-173 | Biolegend | 126522 |
| BV510 Rat Anti-Mouse CD138 (Syndecan-1), Clone 281-2 | Biolegend | 142521 |
| BV510 Rat Anti-Mouse CD45, Clone 30-F11 | Biolegend | 103138 |
| BV510 Rat Anti-Mouse Ig, κ light chain, Clone 187.1 | BD | 742834 |
| BV605 Armenian Hamster Anti-Mouse CD95 (Fas), Clone Jo2 | BD Horizon | 740367 |
| BV650 Rat Anti-Mouse CD93, Clone AA4.1 | BD | 563807 |
| BV650 Rat Anti-Mouse IgG3, Clone R40-82 | BD | 744136 |
| BV711 Rat Anti-Mouse CD138 (Syndecan), Clone 281-2 | Biolegend | 142519 |
| BV785 Rat Anti-Mouse CD138 (Syndecan-1), Clone 281-2 | Biolegend | 142534 |
| BV786 Rat Anti-Mouse IgD, Clone 11-26c.2a | BD Horizon | 563618 |
| FITC Goat Anti-Mouse IgG2c (heavy chain), polyclonal | Abcam | ab97254 |
| FITC Mouse Anti-Mouse CD95 (Fas), Clone SA367H8 | Biolegend | 152606 |
| FITC Rat Anti-Mouse CD45, Clone 30-F11 | Biolegend | 103108 |
| FITC Rat Anti-Mouse/Human GL7 Antigen, Clone GL7 | Biolegend | 144604 |
| Pacific Blue Rta Anti-Mouse/Human GL7, clone GL7 | Biolegend | 144614 |
| PE Mouse Anti-Human IgA, clone IS11-8E10 | Milteny | 130-114-002 |
| PE Rat Anti-Mouse CD138 (Syndecan-1), Clone 281-2 | Biolegend | 142504 |
| PE Rat Anti-Mouse CD98 (4F2), clone RL388 | Biolegend | 128208 |
| PE Rat Anti-Mouse IgA, clone mA-6E1 | eBioscience | 12-4204-82 |
| PE Rat Anti-Mouse IgG2b, Clone RMG2b-1 | Biolegend | 406708 |
| PE Mouse Anti-Mouse/Human Btk (pY223), Clone N35-86 | BD | 562753 |
| PE/Cyanine7 Rat Anti-Mouse IgD, Clone 29-2L17 | Biolegend | 405720 |
| PE/Cyanine7 Rat Anti-Mouse CD39 Antibody, clone Duha59 | Biolegend | 143805 |
| PE/Cyanine7 Rat Anti-Mouse CD98 (4F2), clone RL388 | Biolegend | 128214 |
| PE/Cyanine7 Rat Anti-Mouse IgG2b, clone RMG2b-1 | Biolegend | 406713 |
| PE/Dazzle594 Hamster Anti-MouseCD196 (CCR6), Clone29-2L17 | Biolegend | 129822 |
| PerCP/Cyanine5.5 Rat Anti-Mouse IgM, Clone RMM-1 | Biolegend | 406512 |
| PerCP/Cyanine5.5 Rat Anti-Mouse/Human GL7, clone GL7 | Biolegend | 144610 |
| V450 Rat Anti-Mouse IgG1, Clone A85-1 | BD | 562107 |
| Alexa Fluor 555 Goat Anti-Rabbit IgG (H+L) | ThermoFisher | A-21428 |
| Rabbit anti RFP, Polyclonal | ThermoFisher | 600-401-379 |
| Biotinylated Goat Anti-Mouse IgA, polyclonal | Southern BT | 1040-08 |
| Biotinylated Goat Anti-Mouse IgG, polyclonal | Southern BT | 1030-08 |
| Biotinylated Goat Anti-Mouse IgG2c, polyclonal | Southern BT | 1079-08 |
| Biotinylated Goat Anti-Mouse IgM, polyclonal | Southern BT | 1020-08 |
| Goat F(ab')<sub>2</sub> Anti-Mouse Kappa-UNLB | Southern BT | 1052-01 |
| APC/Cyanine7 Streptavidin | Biolegend | 405208 |

### Immunohistochemistry

Lungs were fixed in 4% PFA for 6 hours at 4°C. Small and large intestines were opened longitudinally, washed with PBS and fixed with PFA 4% for 30 min at 4°C. Afterwards, intestines were washed with PBS and swiss rolls were made and fixed again for another 30 mins. After washing, lungs, lymph nodes and intestines were washed with PBS, incubated overnight in PBS 30% sucrose solution, immersed in OCT and snap-frozen in liquid nitrogen-cooled isopentane. Cryostat sections (10 to 20 μm thick) were dried in silica beads, permeabilized with PBS Saponin 0.5% for 30 minutes and blocked with PBS 0.5% saponin 2% BSA 1% goat serum 1% FCS for 30 minutes. Sections were then incubated with primary antibodies in PBS 0.5% saponin 2% BSA 1% goat serum 1% FCS for at least 1 hour, washed and incubated with secondary antibodies for a further hour. After a final wash, sections were mounted in Fluoromount-G mounting media. Imaging was carried out on a LSM 780 (Zeiss) inverted confocal microscope using a Plan-Apochromat 40x NA 1.3 oil immersion objective or a Plan-Apochromat 20X/0.8 M27 objective in the case of full organ section.

| Antibody | Source | Catalog |
| --- | --- | --- |
| Alexa Fluor 647 Rat Anti-Mouse Bcl-6, clone K112-91 | BD Horizon | 561525 |
| Pacific Blue Rat Anti-Mouse/Human CD45R (B220), Clone RA3-6B2 | Biolegend | 103227 |
| PE Rat Anti-Mouse CD138 (Syndecan-1), Clone 281-2 | Biolegend | 142504 |
| Chicken Anti-GFP, polyclonal | Abcam | ab13970 |

### Analysis of confocal images

After collecting tile scans by confocal microscopy, lung sections were analyzed using Imaris 9.6.0 software. The Surface Creation Wizard was used to identify the YFP positively stained cells based on fluorescence intensity, size and morphology. The selection parameters were established as follows: source channel set to yellow (YFP), smooth: 2, thresholding - background subtraction: 5, threshold: 7, split: 1 and area: 50 μm. Once identified, a value-based visual surface was generated for all positively stained cells, which enabled quantification of fluorescence intensity as well as the frequency of labeled and unlabeled cells. Channel statistics for B220, Bcl-6 and CD138 were obtained for cellular surfaces defined based on YFP+ cells and exported into Excel (Microsoft). Mean voxel fluorescence was plotted in FlowJo software by using the Text to FCS conversion utility (TreeStar Inc). Percentages of YFP+ for B220, GL7 or CD138 were gated using traditional log-scale based flow cytometry plots in FlowJo and then graphed on linear XY plots to map their respective positions within the lung.

### ELISPOT

To measure specific antibody-secreting cells, enzyme-linked immunosorbent spot (ELISpot) multiscreen filtration plates (Millipore) were activated with absolute ethanol, washed with PBS and coated overnight at 4°C with 1μg/ml of nucleoprotein or hemagglutinin (Sino Biological) or 5 μg/ml of Salmonella- or Klebsiella-LPS (Sigma-Aldrich) diluted in PBS. Plates were subsequently blocked for 1 hour with complete medium and incubated for 24 hours at 37°C with serial dilutions of lung, bone-marrow, small and large intestine single cell suspensions. Plates were washed with PBS 0.01% Tween-20 and incubated for 1 hour with 1μg/ml of biotinylated anti-IgM, IgG, IgA or total Ig (Southern Biotech) diluted in PBS 1% BSA. Then, plates were washed and incubated for 30 minutes with 1μg/ml Streptavidin-Alkaline Phosphatase (Sigma). Finally, plates were washed and developed with BCIP®/NBT (Sigma).

### Fecal extract

Fecal pellets were collected at indicated times, snap-frozen and stored at −80°C until analysis. Fecal pellets were resuspended in PBS 10% FCS at a ratio of 10 mg per 100 µl, vortexed for 20 min to disrupt pellets at 4°C and centrifuged 10 min at 13,000 x g. Supernatants were collected and stored at -20°C for subsequent analysis.

### ELISA

To measure the production of specific antibodies, ELISA (enzyme-linked immunosorbent assay) plates were coated overnight at 4°C with either 1μg/ml of hemagglutinin (Sino Biological) or 5 μg/ml of Salmonella LPS (Sigma) diluted in PBS. Plates were washed with PBS 0.01% Tween, blocked for 1 hour at room temperature with PBS 1% BSA and incubated overnight at 4°C with serial dilutions of serum or fecal extract. The following day, plates were washed and probed at room temperature for 1 hour with 1μg/ml of biotinylated anti-IgA or -IgG (Southern Biotech) in PBS 1%BSA. Plates were washed and incubated at room temperature for 30 minutes with 1μg/ml Streptavidin-Alkaline Phosphatase (Sigma). Plates were washed and developed with p-NitrophenylPhosphate (Sigma). 405 nm absorbance was detected using a SPECTRAmax190 plate reader (Molecular Device).

### Gene-expression analysis

Total RNA was extracted using the RNeasy® Mini Kit (Qiagen, Hilden). The reverse transcription into the cDNA was performed with SuperScript II (Invitrogen). The expression of Tgfb1 relative to Actb was measured by real-time PCR (95°C 3 m 1 cycle; 95°C 5 s; 60°C 34 s, 40 cycles, 95°C 15 s, 50°C 1 m 1 cycle) with the StepOnePlus (Applied Biosystems) using ONEGreen® Fast qPCR Premix and the following primers:

Tgfb1 (Fw: 5′-CACAGCTCACGGCACCGGAGA-3′; Rv: 5′-GCTGTACTGTGTGTCCAGGCTCC-3′)

Actb (Fw: 5′-CTCCTGAGCGCAAGTACTCTGTG-3′; Rv: 5′-TAAAACGCAGCTCAGTAACAGTCC-3′).

### A20 class switching and Igha tail deletion

A20 B cells were grown in complete RPMI medium. Guide RNAs (gRNA) were used to induce IgG2a to IgA class switching. Sγ2a 3′ gRNA was designed to target the region between the switch Sγ2a sequence and the constant region Cγ2a sequence. Sα3′ gRNA was designed to target between the switch Sα sequence and the constant region Cα sequence. (Sγ2a 3’gRNA: 5’-TGGCAACATTAGCGCTATCA-3’; Sα3′ gRNA: 5’-GGCTGGAATTAGGCGAAACT-3’). 22 pmoles of Sγ2a 3′ gRNA were coupled to 18 pmoles Cas9-GFP (IDT) to form ribonucleoprotein (RNP1). In the same proportions, Cas9-RFP (IDT) was folded with Sα3′ gRNA to form RNP2. RNP1 was mixed with RNP2 in 1:1 ratio and then added to 2x10^6^ A20 cells, resuspended in an electroporation buffer contained in Cell Line Nucleofector Kit V (Lonza). A20 cells were electroporated (Amaxa Nucleofector II/2b, Program L-013) and sorted one to three days later in a BD FACSAria II (BD, Bioscience). IgA^+^ A20 cells were screened for BCR expression by using goat anti-mouse IgG2a (ThermoFisher Scientific, 10μg/mL) and anti-mouse IgA (mA-6E1, eBioscience, 1μg/mL).

Control Igha wild type tail and Igha deleted tail were obtained by using one gRNA targeting endogenous Igha tail (5’-AGGCCCGTTTGGCAGCAAAG-3’) and a double strand donor DNA (TWIST Bioscience) to allow homologous recombination and selection. The donor DNA contains 5’ to 3’: 5’ homology arm/wild type or deleted Igha tail/three amino-acid linker/P2A sequence/cd90.1 cDNA/3’ homology arm. Homozygous recombinants were screened on 5’ and 3’ side by PCR using the following primers:

5’ Fw: 5’-TCAAGAACTGCTGGGATTCCAGTCCCT-3’

5’ Rv1: 5’-CATCCTGGAGTTGGGTTATGTCTTCCCTTT-3’

5’ Rv2: 5’-AGGCTGTCAGGCTGGTCACCTTCTGCC-3’

3’ Fw1: 5’-TCAAGAACTGCTGGGATTCCAGTCCCT-3’

3’ Fw2: 5’-CATCCTGGAGTTGGGTTATGTCTTCCCTTT-3’

3’ Rv: 5’-GAGTTTGTGGTTTTGCATCAATTTATT-3’

### Phospho-flow cytometry

2 × 10⁶ cells were stained with the LIVE/DEAD Fixable Blue Dead Cell Stain Kit (Thermo Fisher Scientific) for 20 minutes on ice. The cells were plated in 100 μL of RPMI with 2% FCS for 1 hour at 4°C and then stimulated at 37°C with 10 μg/mL Goat anti-mouse IgA (SouthernBiotech) in 100 μL of RPMI with 2% FCS. Cells were fixed for 10 minutes at 37°C and 20 minutes on ice with 200 μL of Fixation/Permeabilization solution (FoxP3/Transcription Factor Fixation and Permeabilization buffer, eBioscience, Invitrogen). Cells were washed with 150 μL of Permeabilization buffer and incubated with 25 μL of hybridoma supernatant 2.4G2, diluted 1:4 in Permeabilization buffer, for 15 minutes on ice. Then, the cells were incubated for an additional 30 minutes on ice with the indicated anti-mouse antibodies. Cells were washed twice and incubated with PE Mouse anti-Btk (pY223) (N35-86, BD Phosflow, 1 μg/mL) or anti-Syk (pY348) (558529, BD Phosflow, 1 μg/mL) for 30 minutes at room temperature in the dark, washed with PBS containing 2% FCS, and analyzed using an LSRIIUV cytometer

### Calcium measurement

8 × 10⁶ cells were loaded with 5 μM Indo-1 (Invitrogen) in RPMI with 10 mM HEPES (pH 7) for 30 minutes at 37°C. The cells were then incubated with RPMI containing 10 mM HEPES (pH 7.4) and 5% FCS in a 1:1 proportion at 37°C for 30 minutes, followed by a wash with RPMI containing 5% FCS. Calcium flux was measured using an LSRIIUV cytometer. The baseline (bound/unbound) was acquired for 1 minute, after which the cells were stimulated with anti-IgA (SouthernBiotech) at indicated concentrations, and calcium flux was acquired for 5 minutes. Ionomycin (Sigma-Aldrich) was then added at a concentration of 5 μg/mL, and calcium flux was measured for an additional 5 minutes.

### Western blot

10 × 10⁶ cells were plated in 100 μL of RPMI for 1 hour at 4°C and then stimulated at 37°C with 40 μg/mL Goat anti-mouse IgA (SouthernBiotech) in 100 μL of RPMI. Cells were fixed during 30 minutes on ice with 400µL of 2X lysis buffer (Tris 100mM pH 7.4, EDTA 1mM, 1% n-dodecyl-β-D-maltoside, NaCl 274mM, 20% glycerol), supplemented with anti-protease and anti-phosphatases inhibitors (Leupeptin 20µg/mL, Aprotinin 20µg/mL, PMSF 20µg/mL, TPCK 20µg/mL, LECK 20µg/mL, Na3VO4 2mM, NaF 100mM). After max speed centrifugation during 15 minutes, the supernatants containing the proteins were collected and proteins levels were quantified using a Bradford assay. 80µg of protein were incubated with 10% β-mercaptoethanol and boiled for 5mn at 95°C. Samples were then loaded in the gel and migrated for 4 hours at 150V. Proteins were then transferred for 2 hours on a membrane at 40V. The membrane was blocked overnight with BSA 5% and stained with Anti-Phosphotyrosine Antibody, clone 4G10 (05-321, Sigma, 0.5µg/mL in BSA 5%) for 3 hours. After washes, the membrane was incubated with a Peroxidase AffiniPure Donkey Anti-Mouse IgG (H+L) antibody (JIR715-035-151, Ozyme, 1:5000 in BSA 5%) for 1 hour. After washes, the membrane was developed with Pierce™ ECL Western Blotting Substrate (32106, ThermoFisher) and pictures were acquired thanks to the the Chemiluminescent Western Blot Imager Azure 300 (Azure Biosystems).

